# Enterococcal PrgA extends far outside the cell and provides surface exclusion to protect against unwanted conjugation

**DOI:** 10.1101/2020.06.18.156562

**Authors:** Andreas Schmitt, Helmut Hirt, Michael A. Järvå, Wei-Sheng Sun, Josy ter Beek, Gary M. Dunny, Ronnie P-A Berntsson

## Abstract

Horizontal gene transfer between Gram-positive bacteria leads to a rapid spread of virulence factors and antibiotic resistance. This transfer is often facilitated via Type 4 Secretion Systems (T4SS), which frequently are encoded on conjugative plasmids. However, donor cells that already contain a particular conjugative plasmid resist acquisition of a 2^nd^ copy of said plasmid. They utilize different mechanisms, including surface exclusion for this purpose. *Enterococcus faecalis* PrgA, encoded by the conjugative plasmid pCF10, is a surface protein that has been implicated to play a role in both virulence and surface exclusion, but the mechanism by which this is achieved has not been fully explained. Here, we report the structure of full-length PrgA, which shows that PrgA protrudes far out from the cell wall (approximately 40 nm), where it presents a protease domain. *In vivo* experiments show that PrgA provides a physical barrier to cellular adhesion, thereby reducing cellular aggregation. This function of PrgA contributes to surface exclusion, reducing the uptake of its cognate plasmid by approximately one order of magnitude. Using variants of PrgA with mutations in the catalytic site we show that the surface exclusion effect is dependent on the activity of the protease domain of PrgA. *In silico* analysis suggest that PrgA can interact with another enterococcal adhesin, PrgB, and that these two proteins have co-evolved. PrgB is a strong virulence factor, and PrgA is involved in post-translational processing of PrgB. Finally, competition mating experiments show that PrgA provides a significant fitness advantage to plasmid-carrying cells.

## INTRODUCTION

Infections caused by multi-drug resistant bacteria have become increasingly problematic all over the world. Enterococci are normally considered commensal bacteria and reside in the gastrointestinal tract of humans and other mammals. However, they are also among the leading nosocomial pathogens, most often associated with bloodstream and urinary tract infections [1–3]. Of special concern are *Enterococcus faecalis* and *Enterococcus faecium,* as these two species are known to have highly transmissible mobile genetic elements (MGEs) in their genomes. These MGEs encode various virulence factors as well as resistance to many antibiotics. Enterococci can very efficiently spread these MGEs to other bacteria, including other medically relevant species such as streptococci and staphylococci. This horizontal gene transfer is mostly carried out via Type 4 Secretion Systems (T4SS) that are encoded on these MGEs [4–7]. Enterococci are therefore acknowledged as important reservoirs for the spread of antibiotic resistance.

The conjugative plasmid pCF10 from *Enterococcus faecalis* has for several decades served as an important model to understand the regulation, structural and functional features of T4SS in G+ species [8,9]. pCF10 contains a tetracycline resistance gene and is effectively transferred from donor to recipient cells following induction of the *prgQ* operon by a peptide pheromone secreted by recipient cells. The 24 genes in this operon encode for all proteins needed to process the pCF10 plasmid (67.7 kb), establish a channel between mating cells and subsequently transfer the DNA into the recipient cell. The first module of the operon encodes three surface proteins, PrgA, PrgB and PrgC as well as a small predicted RNA-binding protein called PrgU that is involved in mitigating the toxicity cells associated with production of PrgB [9,10]. All three surface proteins have a characteristic C-proximal sortase-dependent cell wall anchoring motif (LPxTG).

PrgA (also known as Sec10), which is the focus of this study, is a 92 kDa large protein in its mature form. It is one of the first identified pCF10-encoded proteins and has earlier been indicated to be involved in surface exclusion [11,12], which functions to block efficient transfer of a plasmid into a host cell that already carries the same plasmid [13]. Recent work has shown that PrgA also increases biofilm formation upon overexpression, and that it is an important virulence factor in a *Caenorhabditis elegans* infection model [14]. PrgB (also known as Aggregation Substance, or AS) has been extensively characterized, and mediates binding to host cells and tissues [9], the cell wall of recipient bacteria and/or extracellular DNA which leads to biofilm formation [14–17]. Expression of PrgB, after induction of the pCF10 plasmid, has shown to yield a ca 140 kDa fulllength version along with a truncation of about 78 kDa. The ratio between the two variants seems to depend on how strongly PrgB expression is induced as well as in which bacteria it is produced [18]. The amount of the 78 kDa variant is strongly diminished when PrgB is expressed in a *ΔprgA* background in *E. faecalis* [14], implicating that PrgA might be involved in post-translational processing of PrgB.

Although the role of PrgA in surface exclusion was suggested over 30 years ago, little direct experimental evidence has been published. Here we present new data that confirm the role of PrgA in surface exclusion. Furthermore, we report the crystal structure of PrgA, whose structural features show that it presents a protease domain far outside the cellular envelope. This protease domain is shown to play a direct role in surface exclusion. We also show that PrgA plays a direct role in the processing of PrgB, and that its expression contributes to competitive fitness of *E. faecalis.*

## RESULTS

### Protein purification and crystallization

PrgA_28-814_ (lacking the N-terminal signal sequence and C-proximal LPXTG motif) was initially used for protein expression and purification. However, this variant was very prone to degradation during expression and purification, resulting in a ladder of bands on SDS-PAGE after immobilized metal affinity chromatography and size exclusion chromatography (Fig. S1). Six of the most prominent bands were analyzed by mass spectrometry and all represented PrgA fragments (Table S1). Based on the analysis of the mass spectrometry data and bioinformatic analysis, we expressed and purified PrgA_291-541_, which was more resistant to degradation and subsequently used for crystallization experiments.

PrgA_291-541_ crystallized in space group P21 with two molecules in the asymmetric unit. The crystallographic phase problem was solved by means of single-wavelength anomalous dispersion (SAD) using crystals soaked with an iodine compound (Methods), and the crystal structure was refined at a resolution of 1.5 Å.

### PrgA contains a CAP domain

PrgA_291-541_ contains a central alpha-beta-alpha sandwich, which is formed by a three stranded antiparallel β-sheet (residues 435-441 and 487-509) flanked by a layer of α-helices on each side (Fig. 1a). One side contains 3 helices (formed by residues 362-383, 451-468 and 476-482) while the other side harbors two (formed by residues 392-404 and 414-424). The search for homologous proteins based on the amino acid sequence was inconclusive, but a structure based homology search using the DALI server identified significant similarity to proteins from the superfamily of cysteine-rich secretory proteins, antigen 5, and pathogenesis-related 1 proteins (CAP, Pfam: PF00188) [19].

**Figure 1.**
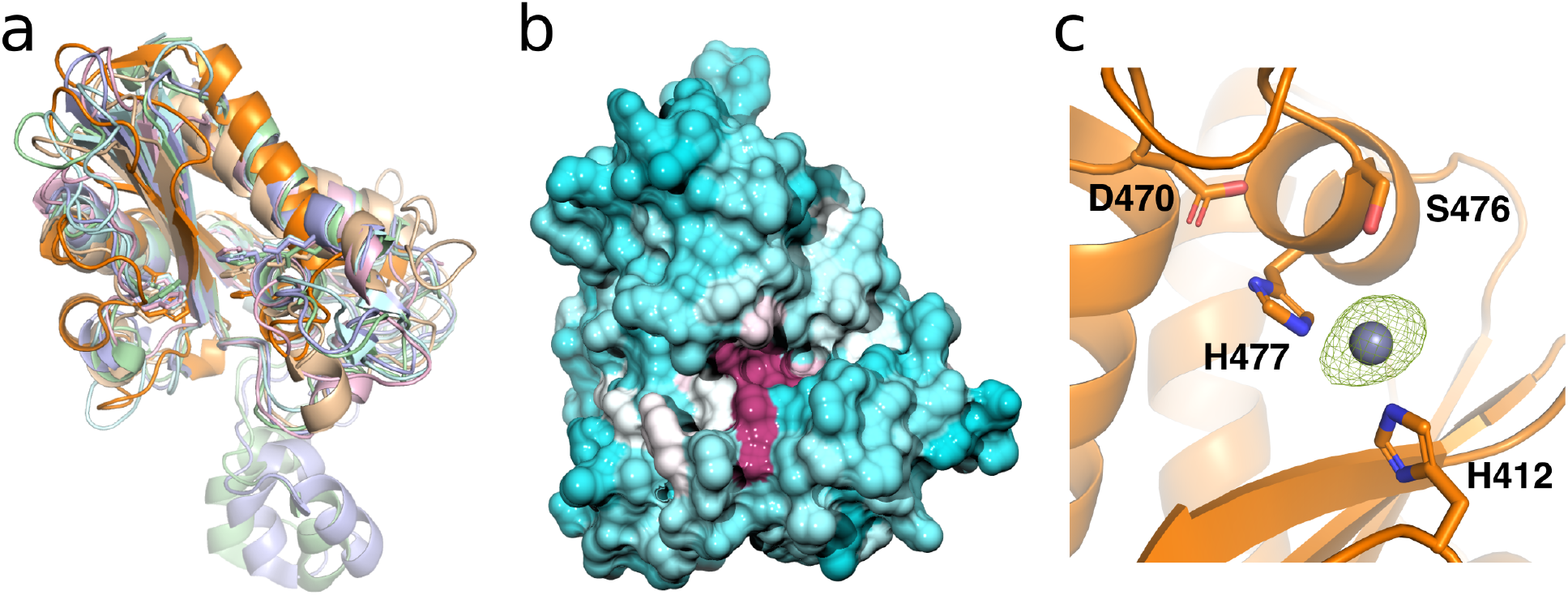
Structure of the CAP domain of PrgA. a) Structural comparison of PrgA (orange) with several other CAP domain proteins available in the PDB (PBB code 1U53: beige, 1XTA: green, 2DDA: blue, 3Q2R: cyan, 5JYS, light purple). Their sequence homology of the CAP domain is low (ca 15%), but they all share the same overall fold. b) Surface representation of the PrgA CAP domain, with the homology to other CAP domains indicated by color, ranging from cyan (no homology), via white, to purple (indicating 100% conserved residues). c) The proposed catalytic triad of the CAP domain in PrgA consists of S476, D470, His412 and His477. A Zn^2+^ atom (gray sphere) was modelled in between the two histidines based on the anomalous difference density collected at the Zn edge (green mesh).

Proteins of this family can be found in a wide range of organisms, spanning from prokaryotes to higher eukaryotes. They have been associated with a plethora of diverse, functions including but not restricted to fungal virulence, ion channel regulation and cell-cell adhesion [20–24].

Sequence comparison of different CAP proteins with PrgA (Fig. S2) revealed, that despite very high structural similarity (Fig. 1a), PrgA lacks all conserved CAP sequence motifs and this explains why this domain was not identified by bioinformatic approaches [22,24]. According to structural superimposition only four residues of PrgA is conserved among known CAP proteins, namely Histidine 412, Glutamate 436, Aspartate 470 and Histidine 477, all of which are located around a cleft on the protein surface (Fig. 1b). This structural feature has earlier been described as a CAP cavity tetrad. It can be found in all structurally characterized CAP domains and was suggested to act as an active site [22,24].

In our electron density we noted a feature located in between His477 and His412 that could originate from a metal ion, likely originating from the crystallization condition. Several structures of different CAP proteins have a divalent cation modeled in between the two conserved histidines [24–26]. We therefore tested a range of different metal ions (Zn^2+^, Mg^2+^, Ca^2+^, K^+^ and Na^+^) for binding to PrgA, with isothermal titration calorimetry (ITC). The only ion that bound with a near physiologically relevant affinity was Zn^2+^, which bound to PrgA with a K_D_ of ca 8.0 μM (Fig. S3). That Zn^2+^ could indeed bind to the predicted metal binding site was verified by soaking PrgA_291-541_ crystals with ZnSO4 and collecting data at the peak of the Zn absorption edge. The corresponding anomalous signal from a single Zn^2+^ ion was located exactly between His477 and His412 (Fig. 1c).

### PrgA contains a potential hydrolase domain

Interestingly, in the CAP cavity of PrgA the His477, Ser476 and Asp470 are positioned as a classical catalytic triad that strongly resembles the active site of a hydrolase. Fitting with this, the amide nitrogen of Ser467 together with Gly474 form a pocket, resembling the so-called oxyanion hole, which acts in stabilization of hydrolase reaction intermediates. However, while the carboxyl group of Asp470 forms a hydrogen bond to the near nitrogen of His477, the imidazole ring is rotated by 90°. This orientation prevents the nucleophile activation of Ser476, thus rendering the catalytic triad inactive (Fig. 1c). The observed rotamer of the imidazole ring is stabilized by the Zn^2+^, which is coordinated between His412 and His477.

As mentioned previously, we had noted that PrgA_28-814_ was heavily degraded during purification. Based on this, the obtained structural data, and published reports from other CAP proteins, we therefore predicted that the CAP domain of PrgA would be a serine protease, and that its protease activity could be inhibited by divalent cations. To investigate if PrgA was a serine protease with Ser476 as the catalytic residue, a PrgA variant containing a Ser476 to Ala mutation (PrgA_28-814(S476A)_) was designed, overexpressed and purified. PrgA_28-814(S476A)_ could be purified to high homogeneity, without any major degradation products (Fig. S4). The same result could also be achieved when we purified wild-type PrgA (PrgA_28-814_) in the presence of 10 mM divalent cations. This indicates that PrgA indeed can function as a serine protease that depends on S476 and is inhibited by divalent cations.

### PrgA presents its CAP domain far outside the cell wall

With the obtained knowledge that the addition of divalent metal ions blocks degradation, intact PrgA_28-814_ was purified and subjected to crystallization trials. Weakly diffracting crystals were obtained in space group P4_3_22 (to ca 20 Å). The diffraction properties of the crystals were subsequently improved via controlled dehydration by sequential transfer of the crystals into solutions with ever higher PEG400 concentrations, going from 37.5% to 90% in 10 steps. The ultimate resolution was 3.0 Å. We solved the phase problem with the previously solved core domain as a molecular replacement model, and subsequently improved the phases using solvent flattening, which worked very well due to the high solvent content (81%).

The overall structure resembles a head on a stick, similar to a tadpole. The core domain (residues 304-530) forms the head that sits at the top. The N-proximal (residues 55-303) and C-proximal (residues 531-798) domain form the stick; a 2-helix coiled coil that protrudes straight away from the core domain (Fig. 2a). The coiled coil domain extends to a total length of 385 Å, making it to our knowledge the longest 2-helix coiled-coil reported in bacteria to date. Only the first 27 and last 16 residues of the protein construct used for crystallization are missing in the electron density. Since the C-terminus of PrgA is anchored to the cell wall *in vivo,* the long coiled-coil positions the CAP domain ~40 nm away from the cell wall. Two proline residues (P131 and P707) are located next to each other approximately 15 nm from the anchoring point in the cell wall, causing a kink in the coiled coil (Fig. 2b). This kink region provides increased flexibility compared to the otherwise relatively rigid coiled coil.

**Figure 2.**
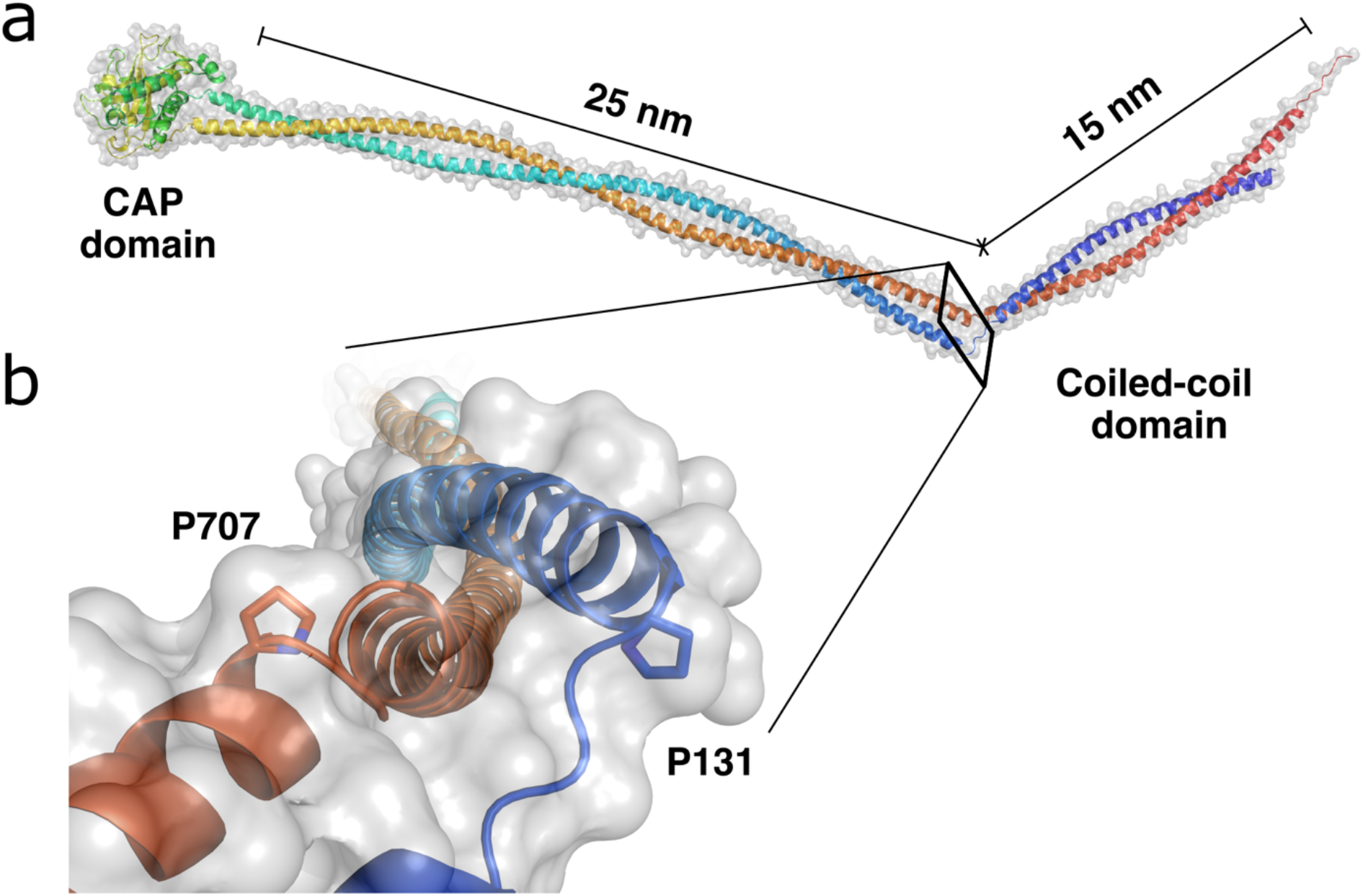
Structure of PrgA_28-814_, with the protein in a cartoon (rainbow colored from N-terminal, blue, to C-terminal, red) representation and its surface as a translucent gray. a) The structure of PrgA consists of a CAP domain (shown on the left), formed by the residues of roughly the middle of the protein sequence, and a coiled-coil domain that is ca 40 nm long. The cell-wall anchoring motif is located at the C-terminus of full-length PrgA, which would mean the CAP domain is presented around 40 nm away from the cell-wall surface. b) Two prolines (P131 and P707), one in each helix, are located close to each other in the coiled-coil domain resulting in a kink.

### Surface expression of PrgA in donor cells reduces aggregation and transfer

The elongated structure of PrgA that we determined by X-ray crystallography, raised the possibility that the protein architecture itself could play a key part in surface exclusion. Since PrgA can protrude 40 nm out from the cell surface, it could spatially interfere with close contacts between cells and in this way prevent mating pair formation and subsequent conjugation. This hypothesis was first tested by comparing the aggregation phenotype of monocultures of cells harboring: 1) wild-type pCF10, 2) pCF10 with the putative active site serine of *prgA* mutated (pCF10:*prgA*::S476A) or 3) pCF10 deleted for *prgA* (pCF10:Δ*prgA*). After induction, cells were pelleted, resuspended in PUM-buffer and vortexed. We found that aggregates reformed significantly faster in the pCF10:Δ*prgA* strain compared to both pCF10 and pCF10:*prgA*:S476A (Fig. 3a & S5). This suggests that the role of PrgA in reduction of cell clumping/aggregation is not influenced by its serine protease activity, but could indeed be a direct result of its extended structure.

**Figure 3.**
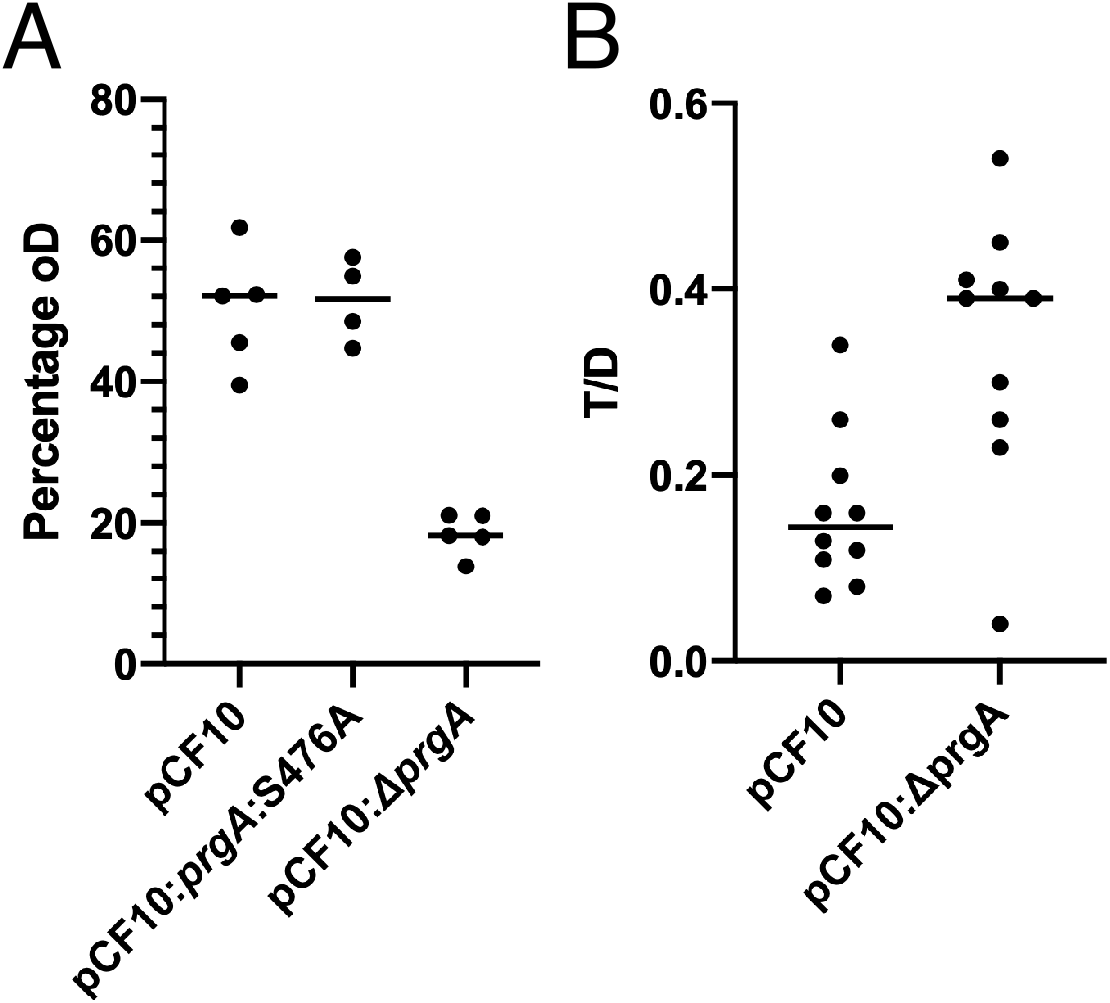
Phenotypes of pCF10:*prgA*::S476A and pCF10:Δ*prgA*. (A) Aggregate formation is increased upon deletion of *prgA,* but unaffected by mutating its putative active site serine. Cells harboring wildtype PrgA (pCF10), PrgA:S476A or pCF10 with an in-frame deletion of *prgA* were induced with cCF10, pelleted, resuspended in PUM-buffer and vortexed. The optical density (OD) of the supernatant was measured as a proxy for the strength of aggregate formation after 15 min of settling. Lower OD indicates stronger aggregate formation (as aggregates sink to the bottom and are not measured). One-way ANOVA gives a p-value of <0.0001, and a p-value of 0.0005 when performing a Welch’s t-test (B) Absence of PrgA increases donor ability. 15 minute matings were performed with either OG1RF:pCF10 or OG1RF:pCF10:Δ*prgA* as donor cells and OG1Sp as recipient cells. Donor cells were induced with 10 ng/ml cCF10 for 45 min before contact with recipient cells. Transconjugants were selected on plates containing tetracycline and spectinomycin and enumerated. T/D: transconjugant/donor. p=0.013, Unpaired t-test with Welch’s correction.

We then investigated if this PrgA dependent interference with the aggregation process also led to a lower donor efficiency. We compared the donor efficiency for cells carrying pCF10 or pCF10:Δ*prgA*. As expected, the lack of PrgA on the cell surface significantly increased the donor efficiency to a plasmid-free recipient (Fig. 3b). Presumably the absence of PrgA on the pCF10:Δ*prgA* donor cells allowed for closer contacts with recipient cells. When this assay was done with the *prgA*:S476A derivative of pCF10, we did not detect a difference in donor ability relative to wild type pCF10 (data not shown), suggesting that the putative protease motif was not required for PrgA-dependent reduction in donor ability shown in Fig. 3B.

### Expression of PrgA on the recipient cell surface reduces pCF10 transfer

If the role of PrgA in surface exclusion is indeed due to spatial interference, we expect the same effect of PrgA expression in the recipient cell. We therefore examined transfer into pCF10- containing recipient cells using donor cells containing the pCF10 derivative pCF10/G2, which encodes a gentamicin resistance marker. As predicted, the recipient cells carrying pCF10:Δ*prgA* had a significantly higher number of transconjugants per donor cell, relative to the wild-type pCF 10 strain (Fig. 4a). There also appeared to be a slight increase in transconjugants in the *prgA*:S476A mutant strain, but the difference from the wild-type strain was not statistically significant (p=0.08).

To examine the role of PrgA expression in recipient cells (independent of other pCF10 gene products), the *prgA, prgA:S476A* and *ΔprgA* alleles were cloned into the cCF10 inducible expression vector pCIE [27]. The expression of wild-type PrgA from pCIE led to a 5-10 fold reduction in transfer of the pCF10/G2 plasmid from donor cells (Fig. 4b). Notably, expression of <u>PrgA</u>:S476A did not provide surface exclusion to the same extent. This suggests that the postulated active site in the CAP-domain of PrgA is required for its full surface exclusion activity (Fig. 4b).

### PrgA expression confers a competitive fitness advantage in mixed cultures

To investigate the effect of PrgA on the competitive fitness of pCF10-carrying strains, OG1Sp:pCF10/G2 and OG1RF:pCF10 (or *prgA* allelic variants) cells were placed on an agar surface at high density and grown for three days in the presence of the plasmid-free strain OG1ES to provide pheromone induction. The data, normalized to the recipient cell numbers, showed that plasmid transfer into pCF10 containing cells from pCF10/G2 donor cells was increased in cells lacking *prgA,* as expected from earlier results (Fig. S6). Recipient cells harboring pCF10:*prgA*::S476A showed an intermediate level of transfer (Fig. S6). Table 1 shows that the ratio of donors with pCF10/G2 (D2) to donors with pCF10 (D1) as percentage of the population in all three experimental set-ups was around 1 at the outset of the experiment. In the pCF10/G2 - pCF10 pairing the D1/D2 ratio only showed a mild increase over three days for wild-type pCF10 (less than 2-fold, from 1.2 to 2.6 p=0.24), but this ratio increased much more in the *prgA*::S476A mutant (ratio increased 9-fold from 0.8 to 7.3, p=0.047) and further shifted in the pCF10:*ΔprgA* mixture (D1/D2 ratio increased 11-fold from 0.7 to 7.7, p=0.020). This data indicates a loss of cell viability or growth disadvantage for cells carrying *prgA*::S476A or *ΔprgA* mutant alleles as compared to wild-type under the conditions of this experiment.

**Table 1.** Comparison of the cell numbers from two donor strains (D1 and D2) in relation to the recipient strain after a three-day competition experiment indicate that active PrgA increases donor cell fitness. This data from the Triple Mating competition experiment depicted in Fig. S6 (the number of cells on day 3) show cell numbers of the population of Donor 1 (D1; OG1Sp:pCF10/G2) and Donor 2 (D2; OG1RF:pCF10 and variants, vertical column). These values are reported in relation to the number of recipient OG1ES cells (Value = 1). The ratio of D1/D2 on Day 0 is compared to the ratio of the two strains on Day 3. The two competitions involving altered D2 (pCF10:*prgA*::S476A and *pCF10:ΔprgA)* show a significant significant increase in the ratio of D1/D2, indicating that donor cells with these *prgA* variants have a lower fitness. Mean and standard deviation are presented. P-values (between D1 and D2) were determined by unpaired t-test with Welch’s correction. Day 3 data are also shown in Fig 3 for D1 and D2.

|  | Day 0 |  |  |  | Day 3 |  |  |  | Day 3/ Day 0 |
| --- | --- | --- | --- | --- | --- | --- | --- | --- | --- |
|  | D1 | D2 | P-value | Ratio D1/D2 | D1 | D2 | P-value | Ratio D1/D2 | Fold increase in D1/D2 ratio |
| pCF10 | 0.57<br>±0.23 | 0.42<br>±0.4 | 0.50 | 1.4 | 0.26<br>±0.36 | 0.065<br>±0.058 | 0.24 | 2.6 | 1.9 |
| pCF10:<br>*prgA*<br>::S476A | 0.66<br>±0.16 | 0.81<br>±0.26 | 0.29 | 0.8 | 0.7<br>±0.49 | 0.086<br>±0.067 | 0.047 | 7.3 | 9.1 |
| pCF10:<br>Δ*prgA* | 0.32<br>±0.19 | 0.48<br>±0.51 | 0.53 | 0.7 | 0.76<br>±0.49 | 0.091<br>±0.089 | 0.020 | 7.7 | 11.0 |

### PrgA is involved in PrgB processing

PrgB can be found in pCF10 containing cells both as a full-length product of 140 kDa as well as in a smaller form of about 78 kDa. The amount of the shorter form is greatly diminished if *prgA* is deleted from pCF10 [14]. Since the data we have described so far indicate that the CAP-domain of PrgA can act as a serine protease, it is therefore tempting to speculate that PrgA is directly responsible for the hydrolysis of PrgB.

The conjugative plasmid pAD1 contains a close homolog of PrgB that is called Aggregation Substance, which has a known cleavage site [28]. This is located just after the adhesion domain, in a predicted unstructured region of PrgB (Fig. S7a). If PrgB would be cleaved there it matches matches with the 78 kDa size of processed PrgB as previously reported [14,18]. To determine if the cleavage site is conserved, we aligned all *prgB* genes in Enterococcus plasmids annotated in the NCBI database. This revealed that the proteolytic site can be divided into 4 main groups (Fig. S7b and Table S2). For PrgB the sequence around the proposed cleavage site is IFNYGNPKEP. To analyse whether PrgA could bind to this sequence, a IFNYGNPKEP peptide was docked onto the structure of the PrgA CAP domain using the CABS-dock webserver in an unbiased manner [29]. The top seven clusters indeed placed the IFNYGNPKEP peptide in the cavity with the active site serine (Fig. 5a).

**Figure 4.**
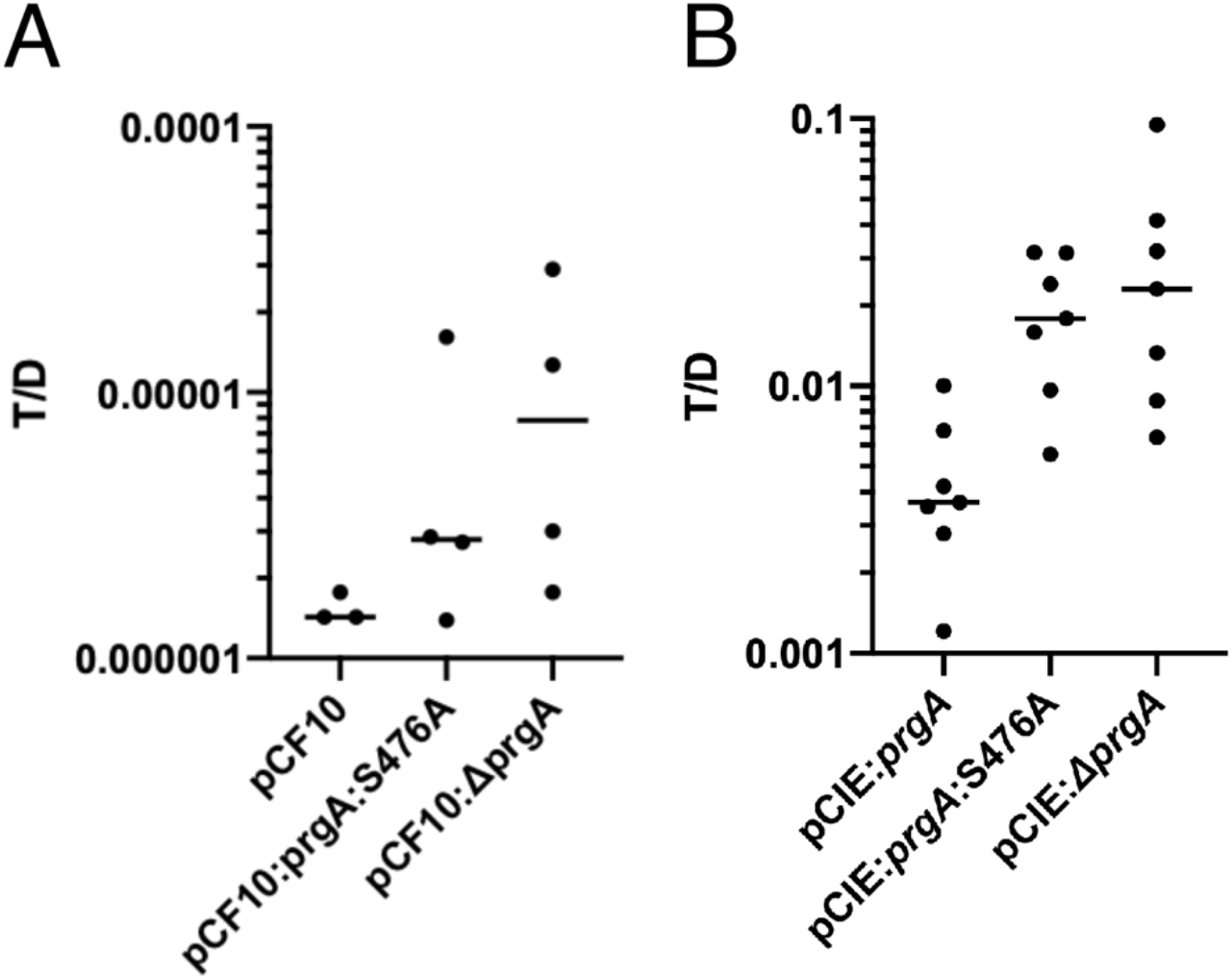
Serine in position 476 is required for effective surface exclusion by PrgA. Matings were performed with a pCF10 derivative as donor and strains containing *prgA*-alleles in a pCF10 (A) or pCIE (B) background. (p-value=0.0489 One-way ANOVA, Mann-Whitney p=0.0041)

We then analysed the sequences of the *prgA* genes from all enterococcal plasmids that also contain *prgB.* This revealed that the *prgA* genes cluster into four distinct groups that coincide with the *prgB* clusters (Table S2 and Fig. S8). Strikingly, when grouping the PrgA homologs based on their sequence of the residues interacting with the PrgB peptide (Fig. 5B), they still group together with the distinct PrgB groups (Table S2). This suggests that the binding cleft in PrgA (containing the predicted active site serine) has thus co-evolved with the sequence around the cleavage site in PrgB, indicating that PrgA might indeed be directly involved in the hydrolysis of PrgB.

**Figure 5.**
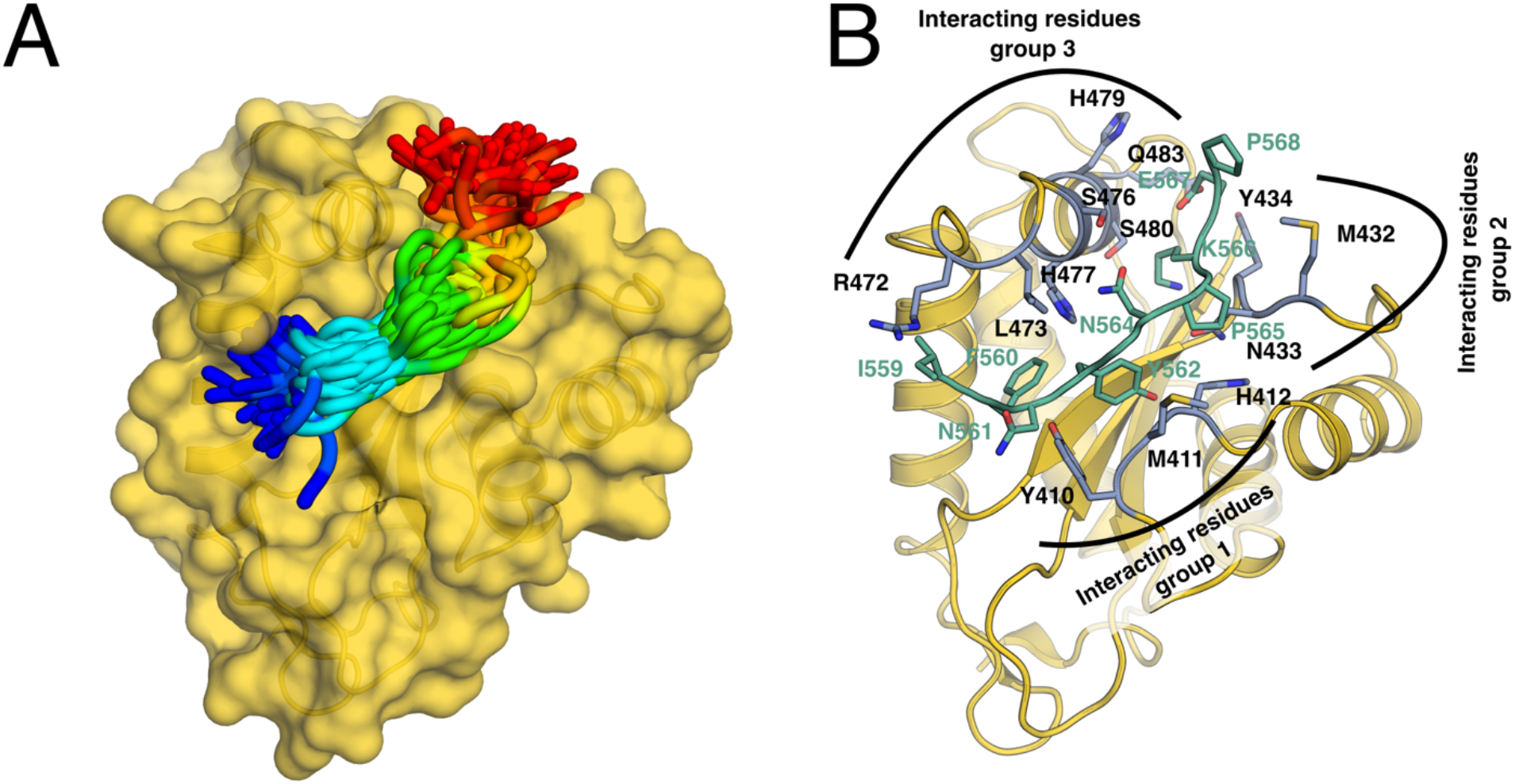
Docking studies of the predicted PrgB proteolysis region onto the structure of the PrgA CAP domain. A: The IFNYGNPKEP peptide (rainbow colored from N-terminal, blue, to C-terminal, red) docked onto the PrgA CAP domain, which is shown in a yellow cartoon and surface representation. B: The CAP domain of PrgA in cartoon representation with potential interacting side chains shown as blue sticks, and a representative docked peptide (with the sequence the predicted proteolysis region in PrgB: IFNYGNPKEP) shown as green sticks.

To test this *in vivo*, Western-Blot analysis (with PrgB-specific antibodies) was done on extracts from induced donor cells with wild-type pCF10, pCF10:Δ*prgA* or pCF10:*prgA*:S476A. In line with previously published findings [14], extracts from wild-type showed a prominent band for the 78 kDa version of PrgB, and a faint band for full-length PrgB, while in the pCF10:Δ*prgA* strain the 78 kDa band was hardly detected and more full-length PrgB was present (Fig. 6). Cells with pCF10:*prgA*:S476A also produced significantly more full-length PrgB, but they also showed considerable amounts of the 78 kDa variant. This suggests that either PrgA:S476A has residual protease activity or that this variant can modulate cleavage of PrgB by another protease. The most prominent proteases in *E. faecalis* are GelE and SprE (Thomas *et al,* 2008). We therefore moved the pCF10 derivatives into both *ΔgelE* and *ΔgelEΔsprE* strain backgrounds to investigate the impact of those proteases on the size of PrgB. However, no effect on the amount of 78 kDa sized PrgB could be observed. *In vitro* proteolysis experiments were also performed, using PrgA_28-814_ with PrgB and BSA as substrates, but no cleavage of either PrgB or BSA could be seen (data not shown).

**Figure 6.**
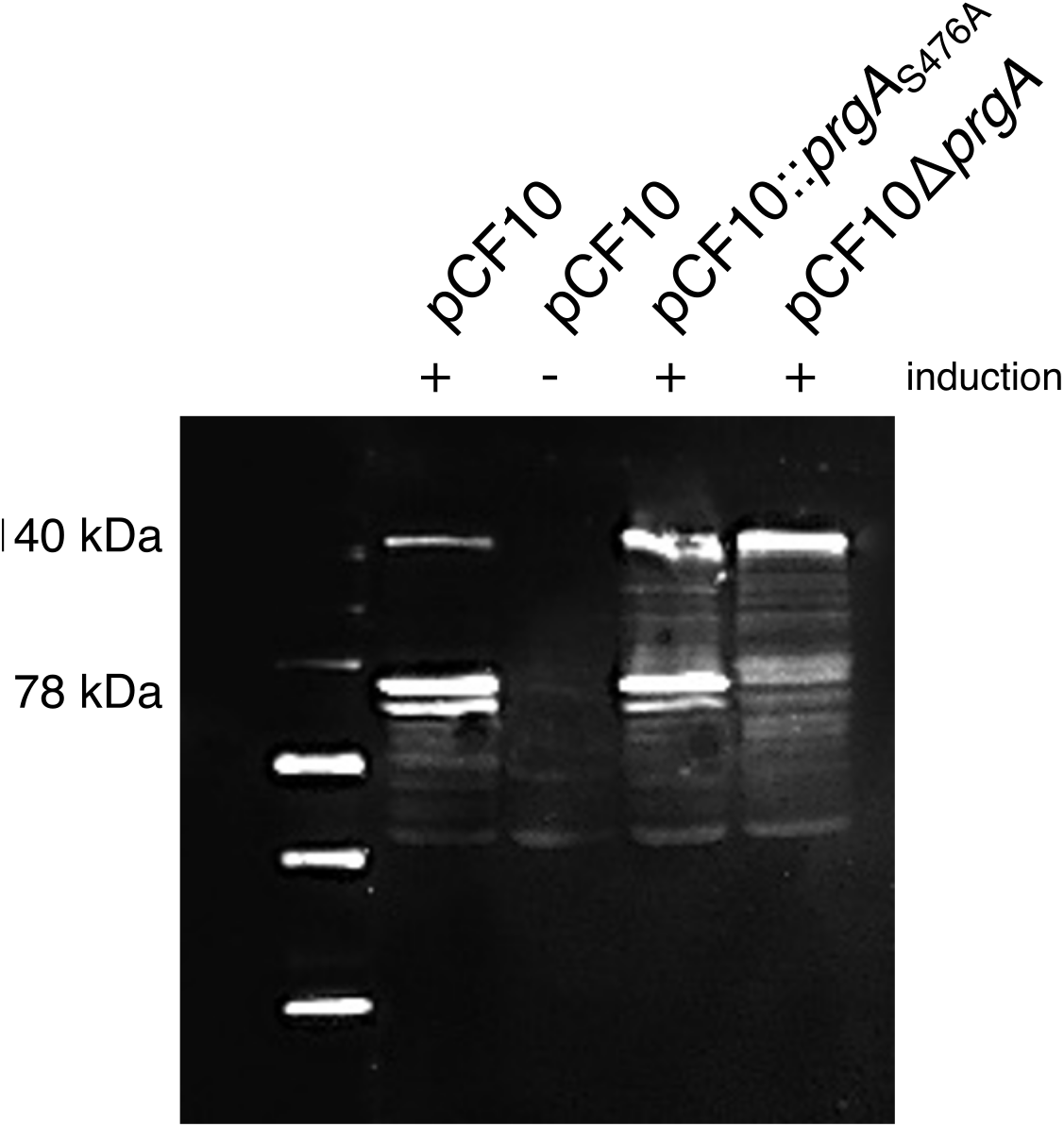
Western-Blot analysis with PrgB-specific antibodies on extracts from cells with induced pCF10 (lane 1), uninduced pCF10 (lane 2), induced pCF10:*prgA*:S476A (lane 3) or induced pCF10:Δ*prgA* (lane 4).

## DISCUSSION

That PrgA plays an important role in surface exclusion in pheromone-responsive plasmids was already established several decades ago. To the best of our current knowledge, all pheromone-responsive plasmids, with the sole exception of the in other aspects unusual plasmid pAM373, encode for a homolog for the pCF10 *prgA* gene. The high degree of homology and conservation indicate the importance of PrgA. More recently, PrgA was found to also play a role during biofilm formation [14]. However, structural information and molecular details about its function have been lacking. The crystal structures of PrgA that we present here, reveal two previously unknown features that could play important roles in its function as a gatekeeper against un-necessary conjugation with other pCF10 containing cells.

First of all, we found that PrgA has a long extended structure that can protrude as far as 40 nm from the cell surface (Fig. 2). This is made possible by a coiled coil domain that extends to a total length of 385 Å, making it the longest 2-helix coiled-coil structure reported from bacteria in the Protein Data Bank to date. Donor cells without PrgA on the surface (pCF10:*dprgA*) aggregated faster and exhibited improved mating efficiency in comparison to wild-type pCF10 cells (Fig. 3). This effect of PrgA (lowering aggregation and mating efficiency) could result from steric hindrance by its long rigid coiled coil domain.

The second significant structural feature that we found is that PrgA has a CAP-domain that is presented ~40 nm away from the cell surface (Fig. 2). This globular domain contains a CAP cavity tetrad consisting of His412, Glu436, Asp470 and His477. Further inspection of the structure revealed that two of these residues (Asp470 and His477) together with Ser476 were positioned in a way that strongly resembles a classical catalytic triad as found in hydrolases (Fig. 1c). In our structure a metal ion is coordinated by the two histidines from the CAP tetrad: His412 and His477. Due to this coordination, the sidechain of His477 is in a confirmation that is not compatible with a hydrolase activity, since it then cannot activate Ser476. Based on the structure we speculated that PrgA might function as a serine-dependent protease that is inhibited by metal ions. Previous literature on CAP domain proteins indicate that this could be the case. One other CAP protein, Tex31 from cone snails, has been indicated to have a Ca^2+^ dependent protease activity that can be blocked by serine protease inhibitors (Milne *et al,* 2003). Another CAP protein (Sav1118, PDB code: 4H0A) has been crystallized with a peptide bound in the proposed active site. However, experimental evidence of proteolytic activity in Tex31 and other proteins is relatively scarce [22,24].

The role of the potential protease activity of PrgA was addressed by exchanging the serine in the putative active site with an alanine (PrgA_S476A_) and studying the effect of this change both *in vivo* and *in vitro.* In contrast to wild-type PrgA where divalent cations needed to be added to all purification buffers to prevent protein degradation, the PrgA_S476A_ variant could be purified as the full-length protein in standard buffers without any metals added. Protease assays were tried *in vitro,* using wild-type PrgA, PrgB and BSA as substrates. However, no activity could be shown in these assays. This might be due to the fact that our *in vitro* conditions don’t reflect the *in vivo* conditions needed for activity. One obvious difference is that both PrgA and PrgB are soluble proteins in these assays, as in contrast to the *in vivo* situation where they are both anchored to the cell wall. To study the role of the putative protease activity of PrgA *in vivo, E. faecalis* recipient cells with a pCIE plasmid containing either *prgA*::S476A, *prgA* or *ΔprgA* were compared in conjugation assays. Surface exclusion in the PrgA::S476A expressing cells was reduced in comparison to the wild-type allele and resembled the *ΔprgA* allele, therefore suggesting that Ser476, and therefore the protease activity of PrgA, has a central role in surface exclusion.

PrgB is the most obvious target for the protease activity of PrgA. It has long been known that induced donor cells contain two major forms of PrgB as visualized by Western-Blots: the 140 kDa full size protein and a band at ca 78 kDa band [14,18]. It was recently discovered that the amount of full-length PrgB is increased and the smaller sized PrgB band cannot be detected in cells with pCF10:Δ*prgA* [14]. The exact cleavage site in the PrgB homologue from pAD1 has been determined and correspond to an unstructured region directly downstream of the adhesin domain [28]. This region is also present in PrgB. We know from previous studies that PrgB is known to form dimers, so the N-terminal fragment could still be present at the cell wall via heterodimeric PrgB, if only one monomer would be cleaved. *In silico* docking indicate that the presumed proteolytic region in PrgB could indeed bind to the active cavity of the CAP domain in PrgA (Fig. 5). Sequence alignments suggest that *prgA* and *prgB* have co-evolved, not only as a whole, but specifically in the regions predicted to be involved in the binding (Fig. S7 & S8, Table S2). The co-evolution of the *prgA* and *prgB* genes could provide a means to specifically prevent mating of a donor cell with a cell containing the same plasmid, while allowing the mating with donor cells with a different plasmid (and therefore another *prgA* / *prgB* pair). This *in silico* data therefore strengthens the suspicion that PrgA binds to the unstructured region of PrgB downstream of the adhesion domain and hydrolyses PrgB at this specific position.

The data up to this point towards that PrgA could be responsible for this cleavage and accomplish its role in surface exclusion by reducing the amount of PrgB and thereby the efficiency of donor-donor cell contact. It is also possible that a PrgB cleavage product could compete with the full-length protein for mediating formation of an effective mating pair. Experimentally we could not detect the cleaved 78 kDa version of PrgB in extracts from cells containing pCF40*dprgA*, while this form is abundant in pCF10 containing cells (also shown in Bhatty *etal,* 20l5). Performing the same analysis with a strain carrying pCF10*prgA*::S476A showed a marked increase in the ratio between the full-length PrgB and 78 kDa fragment, with much more full-length protein observed than in wild-type pCF10. However, the 78 kDa cleaved PrgB fragment could still be detected. That we still see the 78 kDa band in the *prgA*::S476A background could have several explanations. It could be due to that we haven’t found the optimal conditions for the assay, or alternatively that PrgA might not be solely responsible for PrgB processing. When pCF10 and its variant alleles were placed in strains devoid of the prominent *E. faecalis* proteases GelE and SprE no alteration of PrgB degradation activity was seen, thus indicating that neither of those proteases are involved in PrgB processing.

Although PrgA and its presumed protease activity mediated by serine 476 seems to play a significant role in surface exclusion, it is only a minor contributor to the overall exclusion phenotype of pCF10. This is shown by the around 1000-fold stronger surface/entry exclusion of any *prgA* allele exhibited in the pCF10 background, compared to the pCIE plasmid background. This indicates that there is another component involved in the surface/entry exclusion process in pCF10. Based on homologous systems we suggest that this most likely will be on the level of entry exclusion. A similar two factor approach is present in the *Escherichia coli* F-plasmid family. These F-plasmids contain *traS* (surface exclusion) and *traT* (entry exclusion), where TraS provides a ~10-fold contribution and the membrane protein TraT a ~1000-fold reduction to plasmid entry compared to a plasmid-free recipient. An entry exclusion component was recently also identified in the *ICEBs1* element of the Gram-positive *Bacillus subtilis* [30]. The YddJ protein of this element is the sole component that provides entry exclusion (at a level of 1000-fold), but no clear homolog can be found on pCF10. Although the contribution of PrgA in surface exclusion appears to be minor in our plasmid transfer experiments (ca 10-fold reduction), our competition experiment on solid surface indicates that it is a major disadvantage for cells harboring pCF10 to not have functional PrgA under at high cell densities, *e.g.* in biofilms, conducive to formation of potential mating pairs. Both pCF10*dprgA* and pCF10*prgA*::S476A were at a severe disadvantage in comparison to a donor with intact *prgA* after three days of co-culture with a recipient and a competing pCF10 donor strain. These results suggest that formation of spurious mating pairs between plasmid-containing donor cells may confer a high fitness cost whether or not intercellular plasmid transfer actually occurs.

Taken together, our results demonstrate that PrgA has an active role in decreasing cellular adhesion, likely due to steric hindrance. We solved the crystal structure of PrgA and show for the first time that it has a CAP-domain with a hydrolase-like active site, that sticks out ~40 nm outside the cell wall. We also verified that PrgA is active in surface exclusion, as postulated almost 30 years ago, and we now show that this effect is dependent on the active site serine of the CAP-domain of PrgA. *In silico* analysis indicate that a conserved unstructured region of PrgB binds to this active site and gets cleaved, resulting in the 78 kDa PrgB fragment that has been described in literature. Finally, we show that donor cells with PrgA have a competition advantage.

## MATERIALS & METHODS

### Plasmids and cloning

Changes at position 11440 in pCF10 from AGC (Ser) to GCC (Ala) were introduced by PCR. The product was first cloned into pGEMT-easy and verified by sequencing. The initial approach to clone the fragment into pCJK218 was not successful and the fragment was cloned into pCJK47. Transfer into OG1RF:pCF10, integration and excision was successful, detected by MAMA-PCR and confirmed by sequencing, however, due to the use of pCJK47 and the mobilizing plasmid pCF10ΔoriT [31], recombination in the oriT region led to a transfer negative phenotype. The presence of the mutated oriT region was confirmed by restriction digest and PCR. Restoration of the original oriT was accomplished by cloning the WT fragment into pGEMT-easy and obtaining one correct clone in pCJK218. The plasmid was introduced into OG1RF:pCF10::*prgA*:S476A oriT-; integration and excision were successful and a transfer positive plasmid was identified in a plasmid transfer screen. The sequences of *oriT* and *prgA* were confirmed by sequencing. To isolate the *prgA* allele and its variants from the pCF10 background, the alleles were amplified by PCR and cloned into the cCF10 inducible vector pCIE [27].

Plasmid pCF10/G2 is a derivate of pCF10 isolated during transposon library construction of pCF10 using pZXL5 [32] encoding gentamycin resistance. Integration of the transposon occurred on pCF 10 and resulted in the loss of the last 3 amino acids of PrgY. This results in a higher background of plasmid transfer in uninduced cells in a standard mating (Transconjugants/Donor: 10^-4^) compared to uninduced pCF10 cells (Transconjugants/Donor: 10^-6^).

For overexpression and purification purposes, the DNA encoding for PrgA_28-814_, PrgA_291-541_ or PrgA294-754 was amplified from the pCF10 plasmid and cloned into the expression vector p7XC3H using FX cloning [33]. PrgA_28-814_ was also cloned into the pGEX-6P-2 vector to obtained GST- tagged PrgA_28-814_. These constructs don’t contain the C-terminal cell-wall anchoring motif.

### Protein expression and purification

The vectors for overexpression were transformed to *Escherichia coli* BL21(DE3). Cells were grown to an OD_600nm_ of 1.5 in Terrific Broth medium at 37 °C, before lowering the temperature to 16 °C and inducing the expression by the addition of 0.5 mM IPTG. Expression was done overnight (16 h). Cells were harvested and disrupted with a Constant cell disruptor (Constant Systems) at 25 kPsi in 50 mM HEPES/NaOH (pH 7.5), 500 mM NaCl, 30 mM Imidazole (pH 7.8). The lysate was clarified by centrifugation for 30 minutes at 30,000 x g.

The his-tagged protein was purified at 20 °C on Ni-NTA-Sepharose (Macherey-Nagel). The column was washed with 10 column volumes (CV) of 20 mM HEPES/NaOH (pH 7.5), 300 mM NaCl, 30 mM imidazole (pH 7.8) and bound proteins were eluted from the column with the same wash buffer supplemented with 500 mM Imidazole (pH 7.8). GST-tagged proteins were purified on Glutathione-agarose bead (Protino). The beads were washed, and the protein was eluted by addition of 50 mM Hepes pH 7.5, 500 mM NaCl, 5 mM EDTA and 30 mM reduced Glutathione. Both the His-tagged and GST-tagged variants of PrgA were subsequently loaded on a Superdex- 200 16/600 size exclusion chromatography column (GE Healthcare) in 10 mM HEPES/NaOH (pH 7.5) and 200 mM NaCl. Both tags were removed by the addition of HRV 3C protease. Proteins were concentrated to 12 mg/mL using an Amicon Ultra Centrifugal Filter (Merck-Millipore). To purify non-degraded PrgA (without the S476A mutation), 10 mM Mg^2+^ was added before cell lysis and to all subsequent buffers during the purification. Protein purity was assayed via SDS-PAGE. Note that PrgA contains a many positively charged residues, and therefore migrates slower through the gel than expected based on its molecular weight.

### Mass spectrometry

Peptides for mass spectrometry analysis were generated by in-gel digestion of cut out bands indicated in Fig. S1, in the presence of 20 mM ammonium bicarbonate, 0.01% ProteaseMax (Promega) and 6 ng/μl of sequencing grade trypsin (Promega) for 1 h at 50 °C. For analysis by mass spectrometry, the liquid phase of the in- gel digests was acidified using formic acid (final concentration 1%), and insoluble material was removed by centrifugation (20000 x g, 10 min, 4 °C). The in-gel digest samples were analyzed by LC-MS/MS using a HCT Ultra ETDII iontrap mass spectrometer (Bruker) linked to an Acquity-M nano UPLC (Waters). Peptides were separated at a flow rate of 350 nl/min using a 15 cm C18 nano reversed phase column (HSS T3 1.8 μm 75 μm x 150 mm, Waters) and a 15 minutes gradient from 3 to 50 percent acetonitrile in 0.1% formic acid. The MS scans were performed in the enhanced scan mode covering a mass range from 300 to 2200. Acquisition of MS/MS spectra were performed using alternating CID and ETD fragmentation. Processing of the raw data files was performed using Data Analysis software (version 4, Bruker).

### Crystallization and structure determination

Crystals of PrgA_291-541_ were grown at 20 °C by sitting drop vapor diffusion in a condition containing 0.02 M Na/KPO4, 0.1 M Bis-Tris propane pH 6.5, 20% PEG 3350 with a protein concentration of 12 mg/ml and a protein to reservoir ratio of 1:1 in the drop. Crystals of PrgA_28-814_ were grown at 20 °C by sitting drop vapor diffusion in a condition containing 37.5% (v/v) PEG400, 0.1 M Tris/HCl (pH 8.5), 0.114 M LiSO4. The crystals were dehydrated by sequential transfer of the crystals into solutions with ever higher PEG400 concentrations, going from 37.5% to 90% in 10 steps with 20 minutes incubation time per step. Crystals for native data collection were flash cooled in liquid nitrogen. For soaking with iodine, the crystals were transferred into the mother liquor containing 500 mM JBS Magic triangle I3C and were soaked for 30 seconds prior to flash cooling in liquid nitrogen.

X-ray diffraction data were recorded at beamlines ID 29 and ID30-A at the European Synchrotron Radiation Facility (ESRF), Grenoble, France. The data was processed using XDS [34]. The PrgA_291-541_ crystals belonged to the monoclinic space group P12_1_1 and contained two molecules in the asymmetric unit. The crystallographic phase problem was solved by means of single-wavelength anomalous dispersion (SAD) using I3C soaked protein crystals. The iodine sites were found and refined by SHELX and an initial model was built by ARP/wARP [35,36]. The structure of the high-resolution native data was solved by molecular replacement with PHASER [37] using the initial PrgA_291-541_ structure as a search model. The PrgA_28-814_ crystals belonged to space group P4_3_22 and contained one molecule in the asymmetric unit. PrgA_291-541_ was used as a molecular replacement model to solve the phase problem. Building of the models were conducted in COOT [38]. PrgA_28-814_ and PrgA_291-541_ were refined at a resolution of 3.0 and 1.50 Å, respectively, using PHENIX [39]. For complete data collection and refinement statistics see Table 2. For PrgA_28-814_ and PrgA_291-541_ 92.06 and 98.97 %, respectively, of the residues are located in the favored region of the Ramachandran plot as determined by MolProbity [40]. For PrgA_28-814_ the exact register of the lower part of the coiled-coil (residues 55-132 and 700-798) was difficult to place due to the relatively low resolution.

**Table 2.**
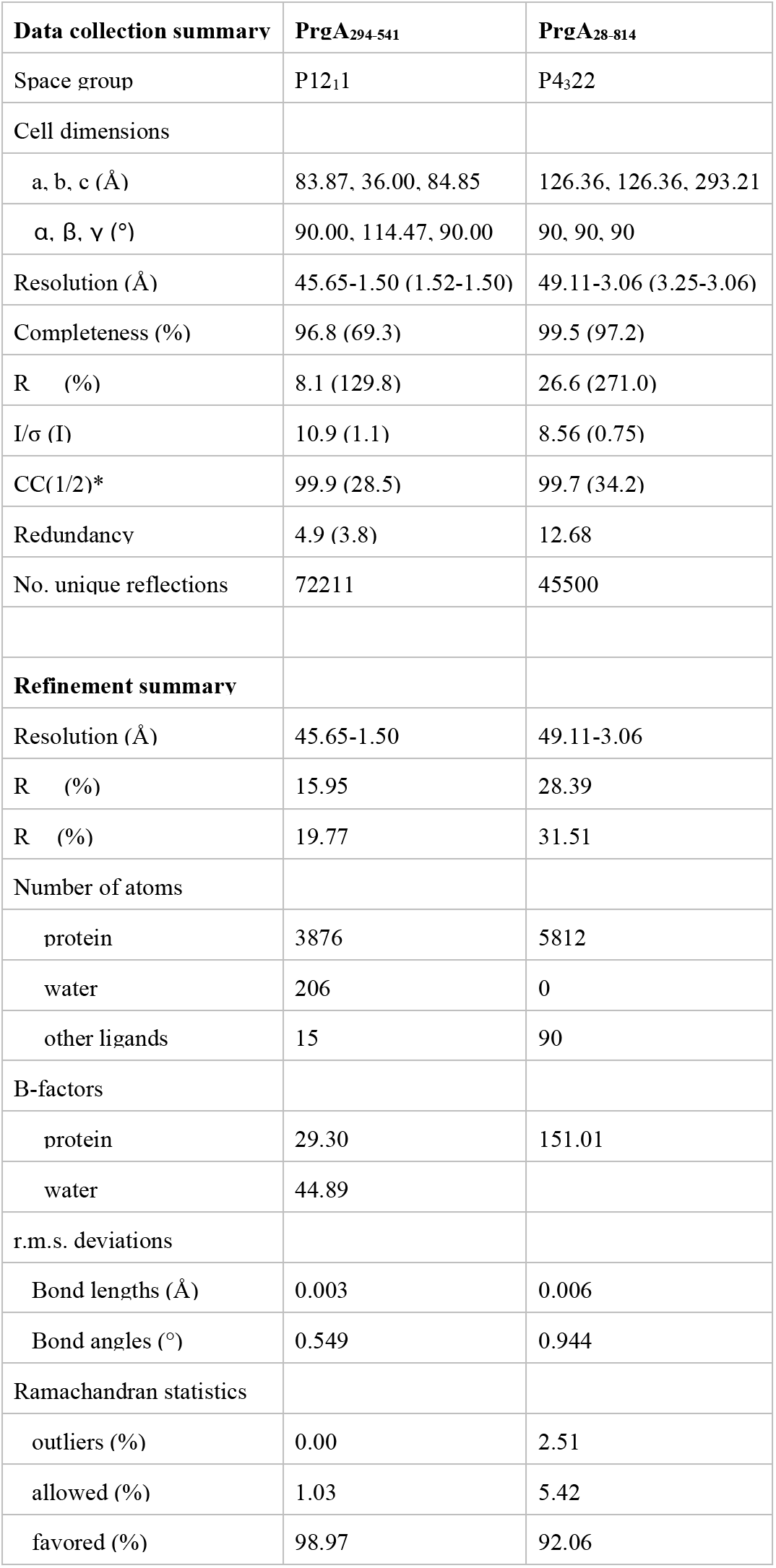
Data collection and refinement statistics.

### Isothermal Titration Calorimetry

The ITC experiments were performed on a MicroCal iTC200 (Malvern). 1mM ZnSO4 solution was titrated into a solution with 0.05 mM PrgA_294-754_. Both solutions were prepared with identical buffer solution, containing 20 mM HEPES pH 7.0 and 150 mM NaCl. The injection sequence consisted of an initial injection of 0.5 μL to prevent artifacts arising from the filling of the syringe (not used in data fitting), followed by injections of 2 μL aliquots with a 120 s interval between each injection. To correct for heats of dilution and mixing, blank titrations of ZnSO_4_ into buffer were subtracted from the ZnSO4-protein titration. The resulting titration curves were fitted to a one-site model using the Origin ITC software package supplied by Malvern to numerically obtain the apparent binding constant (K_D_), the number of binding sites (n) and binding enthalpy ΔH.

### Aggregate Formation Assay

Cells were inoculated 1:100 in Todd-Hewitt Broth (THB) and grown at 37°C for 4 hours. 50 ng/ml cCF10 was added to the induced cultures at time of inoculation. After inoculation, cells were resuspended 10 times with a P1000 tip to homogenize the culture. 1 ml was removed and pelleted at 16000 xg for 5 min in a centrifuge. The pellet was resuspended in 1 ml of PUM-buffer (Rosenberg 1980) and then vortexed at top speed (Genie 2) for 10 seconds. The cultures were allowed to settle for 15 minutes, before 800 μl were removed from the top to measure the oD600. The difference in oD between uninduced and induced cultures served as measure for aggregate formation with lower oD of the supernatant indicating increased aggregate formation.

### Plasmid Transfer Assay (Mating)

Donor (OG1Sp:pCF10/G2) and recipients (OG1RF:pCIE:*prgA*/pCIE:*prgA*:S746A/pCIE:Δ*prgA*) were inoculated 1:10 in THB-Broth and grown separately for one hour. cCF10 was added at 2.5 ng/ml for the donor and 10 ng/ml for the recipients at time of inoculation. After one hour, the recipient culture was inoculated 1:10 with the donor culture and incubated for a further two hours to allow mating to occur. After the completed mating, an aliquot of cells was removed, mixed 1:1 with PUM-buffer, vortexed for 10 seconds and then serially diluted. Donors and transconjugants were enumerated on spectinomycin (1000 μg/ml) plus tetracycline plates (10 μg/ml) and rifampicin (200 μg/ml) plus gentamycin plates (1000 μg/ml), respectively. For the mating shown in the supplemental material, the donor was not induced, recipients were induced with 10 ng/ml cCF10 and mating was only allowed to proceed for 15 min.

### Triple Mating

Population analysis of the competitiveness of the *prgA* mutants was performed on an agar surface created by filling 1 ml of melted THB+1.5% agar in a standard Eppendorf-tube. Overnight cultures were 1:10 diluted in K-PBS and a mating mixture of the three strains was created in a ratio of 1:1:2 of donor 1 (D1) OG1Sp:pCF10-G2, donor 2 (D2) OG1RF:pCF10 (or pCF10:*prgA*::S476A or pCF10:Δ*prgA*) and recipient (R) OG1ES. Cell numbers of the Day 0 mating mixture were immediately determined by serial dilution. 20 μl of the mating mixture was placed on the agar surface in the prepared Eppendorf Tube and incubated for 3 days at 37°C. After 3 days, the cells were removed from the agar surface by resuspension and cell numbers of D1, D2, R and the possible transconjugants (TC1-3) were determined by serial dilution and plating on selective plates. Transconjugants formed from the transfer of pCF10-G2 into D2 (also referred to as Test strain) were labeled TC1. Transconjugants resulting from plasmid transfer into the recipient OG1ES from both D1 and D2 are referred to as TC2. Transconjugants from the transfer of pCF10-G2 into OG1ES recipients are identified as TC3. Selective plates contained 1000 μg/ml spectinomycin, 10 μg/ml tetracycline (D1); 200 μg/ml rifampicin, 10 μg/ml tetracycline (D2); 200 μg/ml rifampicin, 1000 μg/ml gentamycin (TC1), 20 μg/ml erythromycin (R); 20 μg/ml erythromycin, 10 μg/ml tetracycline (TC2) and 20 μg/ml erythromycin and 1000 μg/ml gentamycin (TC3).

### Western Blot

Western Blots were performed using standard procedures. Equal amounts of cell extracts were run on a 7.5% minigel and transferred to a Protran BA85 membrane (Whatman). After overnight blocking, antibody against aggregation substance [41] was incubated using in a 1:2500 dilution for 2 hours. A goat-anti rabbit HRP conjugated secondary antibody (Invitrogen) was incubated using a 1:2500 dilution for 1 hour. The signal was detected by using Supersignal West Pico (Thermo Fisher)

### Peptide docking

The structure of the CAP domain of PrgA and sequence of the region around the proposed PrgB cleavage site was used as input into the CABS-dock server, which was run using default settings [29].

## Accession numbers

Atomic coordinates and structure factors of the PrgA crystal structures have been deposited with the Protein Data Bank (PDB id 6Z9K and 6Z9L).

## Acknowledgements

The authors thank Dr. Eric Geertsma for providing the plasmids of the FXCloning system. We are grateful to the beamline scientists at the ESRF for providing assistance in using beamline ID29 and ID30-A. This work was supported by grants from the Swedish Research Council (2016-03599), Knut and Alice Wallenberg Foundation and Kempestiftelserna (JCK-1524 & SMK-1869) to R.P-A.B., and by PHS grant R35 GM118079 from the National Institutes of Health to G.M.D.

## Author contributions

A.S. performed protein purification, structure determination, *in vitro* protease assays and ITC measurements. H.H. performed cloning and *in vivo assays.* W-S.S. performed PrgB processing assays and western blots. M.J. performed the *in silico* studies. A.S., G.D., J.t.B. and R.P-A.B. planned the experiments, performed the data analysis and wrote the manuscript with input from all authors.

## References

[1] Hidron, A.I. Edwards, J.R. Patel, J. Horan, T.C. Sievert, D.M. Pollock, D.A. Fridkin, S.K. Team, N.H.S.N. & Facilities, P.N.H.S.N., (2008). NHSN annual update: antimicrobial-resistant pathogens associated with healthcare-associated infections: annual summary of data reported to the National Healthcare Safety Network at the Centers for Disease Control and Prevention, 2006-2007., Infect. Control Hosp. Epidemiol. 29 996–1011.

[2] Gilmore, M.S. Lebreton, F. & van Schaik, W., (2013). Genomic transition of enterococci from gut commensals to leading causes of multidrug-resistant hospital infection in the antibiotic era., Curr. Opin. Microbiol. 16 10–16.

[3] Lebreton, F. van Schaik, W. McGuire, A.M. Godfrey, P. Griggs, A. Mazumdar, V. Corander, J. Cheng, L. Saif, S. Young, S. Zeng, Q. Wortman, J. Birren, B. Willems, R.J.L. Earl, A.M. & Gilmore, M.S., (2013). Emergence of epidemic multidrug-resistant Enterococcus faecium from animal and commensal strains., MBio. 4 e00534–13.

[4] Paoletti, C. Foglia, G. Princivalli, M.S. Magi, G. Guaglianone, E. Donelli, G. Pruzzo, C. Biavasco, F. & Facinelli, B., (2007). Co-transfer of vanA and aggregation substance genes from Enterococcus faecalis isolates in intra- and interspecies matings, J. Antimicrob. Chemother. 59 1005–1009.

[5] Arias, C.A. Panesso, D. Singh, K. V Rice, L.B. & Murray, B.E., (2009). Cotransfer of antibiotic resistance genes and a hylEfm-containing virulence plasmid in Enterococcus faecium., Antimicrob. Agents Chemother. 53 4240–4246.

[6] Laverde Gomez, J.A. Hendrickx, A.P.A. Willems, R.J. Top, J. Sava, I. Huebner, J. Witte, W. & Werner, G., (2011). Intra- and interspecies genomic transfer of the Enterococcus faecalis pathogenicity Island, PLoS One. 6 e16720.

[7] Arias, C.A. & Murray, B.E., (2012). The rise of the Enterococcus: beyond vancomycin resistance., Nat. Rev. Microbiol. 10 266–278.

[8] Dunny, G.M. & Berntsson, R.P.-A., (2016). Enterococcal sex pheromones: Evolutionary pathways to complex, two-signal systems, J. Bacteriol. 198. https://doi.org/10.1128/JB.00128-16.

[9] Dunny, G.M., (2013). Enterococcal Sex Pheromones: Signaling, Social Behavior, and Evolution., Annu. Rev. Genet. 47 457–482.

[10] Bhatty, M. Camacho, M.I. González-Rivera, C. Frank, K.L. Dale, J.L. Manias, D.A. Dunny, G.M. & Christie, P.J., (2016). PrgU: A Suppressor of Sex Pheromone Toxicity in Enterococcus faecalis., Mol. Microbiol. 103 398–412.

[11] Dunny, G.M. Zimmerman, D.L. & Tortorello, M.L., (1985). Induction of surface exclusion (entry exclusion) by Streptococcus faecalis sex pheromones: use of monoclonal antibodies to identify an inducible surface antigen involved in the exclusion process., Proc. Natl. Acad. Sci. U. S. A. 82 8582–8586.

[12] Olmsted, S.B. Erlandsen, S.L. Dunny, G.M. & Wells, C.L., (1993). High-resolution visualization by field emission scanning electron microscopy of Enterococcus faecalis surface proteins encoded by the pheromone-inducible conjugative plasmid pCF10., J. Appl. Microbiol. 175 6229–6237.

[13] Garcillán-Barcia, M.P. & de la Cruz, F., (2008). Why is entry exclusion an essential feature of conjugative plasmids?, Plasmid. https://doi.org/10.1016/j.plasmid.2008.03.002.

[14] Bhatty, M. Cruz, M.R. Frank, K.L. Gomez, J.A.L. Andrade, F. Garsin, D.A. Dunny, G.M. Kaplan, H.B. & Christie, P.J., (2015). Enterococcus faecalis pCF10-encoded surface proteins PrgA, PrgB (aggregation substance) and PrgC contribute to plasmid transfer, biofilm formation and virulence., Mol. Microbiol. 95 660–677.

[15] Kohler, V. Keller, W. & Grohmann, E., (2018). Enterococcus adhesin PrgB facilitates type IV secretion by condensation of extracellular DNA., Mol. Microbiol. 109 263–267.

[16] McCormick, J.K. Hirt, H. Dunny, G.M. & Schlievert, P.M., (2000). Pathogenic Mechanisms of Enterococcal Endocarditis., Curr. Infect. Dis. Rep. 2 315–321.

[17] Chuang, O.N. Schlievert, P.M. Wells, C.L. Manias, D.A. Tripp, T.J. & Dunny, G.M., (2009). Multiple functional domains of Enterococcus faecalis aggregation substance Asc10 contribute to endocarditis virulence., Infect. Immun. 77 539–548.

[18] Hirt, H. Erlandsen, S.L. & Dunny, G.M., (2000). Heterologous inducible expression of Enterococcus faecalis pCF10 aggregation substance asc10 in Lactococcus lactis and Streptococcus gordonii contributes to cell hydrophobicity and adhesion to fibrin., J. Appl. Microbiol. 182 2299–2306.

[19] Holm, L. & Laakso, L.M., (2016). Dali server update, Nucleic Acids Res. 44 W351–W355.

[20] Yeats, C. Bentley, S. & Bateman, A., (2003). New knowledge from old: In silico discovery of novel protein domains in Streptomyces coelicolor, BMC Microbiol. 3 1–20.

[21] Guo, M. Teng, M. Niu, L. Liu, Q. Huang, Q. & Hao, Q., (2005). Crystal structure of the cysteine-rich secretory protein stecrisp reveals that the cysteine-rich domain has a K+ channel inhibitor-like fold., J. Biol. Chem. 280 12405–12412.

[22] Gibbs, G.M. Roelants, K. & O’Bryan, M.K., (2008). The CAP superfamily: cysteine-rich secretory proteins, antigen 5, and pathogenesis-related 1 proteins--roles in reproduction, cancer, and immune defense., Endocr. Rev. 29 865–897.

[23] Schneiter, R. & Di Pietro, A., (2013). The CAP protein superfamily: Function in sterol export and fungal virulence, Biomol. Concepts. 4 519–525.

[24] Darwiche, R. Kelleher, A. Hudspeth, E.M. Schneiter, R. & Asojo, O.A., (2016). Structural and functional characterization of the CAP domain of pathogen-related yeast 1 (Pry1) protein, Sci. Rep. 6 28838.

[25] Shikamoto, Y. Suto, K. Yamazaki, Y. Morita, T. & Mizuno, H., (2005). Crystal structure of a CRISP family Ca2+-channel blocker derived from snake venom, J. Mol. Biol. 350 735–743.

[26] Suzuki, N. Yamazaki, Y. Brown, R.L. Fujimoto, Z. Morita, T. & Mizuno, H., (2008). Structures of pseudechetoxin and pseudecin, two snake-venom cysteine-rich secretory proteins that target cyclic nucleotide-gated ion channels: Implications for movement of the C-terminal cysteine-rich domain, Acta Crystallogr. Sect. D Biol. Crystallogr. 64 1034–1042.

[27] Weaver, K.E. Chen, Y. Miiller, E.M. Johnson, J.N. Dangler, A.A. Manias, D.A. Clem, A.M. Schjodt, D.J. & Dunny, G.M., (2017). Examination of Enterococcus faecalis toxinantitoxin system toxin Fst function utilizing a pheromoneinducible expression vector with tight repression and broad dynamic range, J. Bacteriol. 199 e00065–17. https://doi.org/10.1128/JB.00065-17.

[28] Nakayama, J. Watarai, H. Isogai, A. Clewell, D.B. & Suzuki, A., (1992). C-Terminal Identification of AD74, a Proteolytic Product of Enterococcus faecalis Aggregation Substance: Application of Liquid Chromatography/Mass Spectrometry, Biosci. Biotechnol. Biochem. 56 127–131. https://doi.org/10.1271/bbb.56.127.

[29] Kurcinski, M. Jamroz, M. Blaszczyk, M. Kolinski, A. & Kmiecik, S., (2015). CABS-dock web server for the flexible docking of peptides to proteins without prior knowledge of the binding site, Nucleic Acids Res. 43 419–424. https://doi.org/10.1093/nar/gkv456.

[30] Avello, M. Davis, K.P. & Grossman, A.D., (2019). Identification, characterization and benefits of an exclusion system in an integrative and conjugative element of Bacillus subtilis, Mol. Microbiol. 112 1066–1082. https://doi.org/10.1111/mmi.14359.

[31] Staddon, J.H. Bryan, E.M. Manias, D.A. Chen, Y. & Dunny, G.M., (2006). Genetic characterization of the conjugative DNA processing system of enterococcal plasmid pCF10., Plasmid. 56 102–111.

[32] Zhang, X. Paganelli, F.L. Bierschenk, D. Kuipers, A. Bonten, M.J.M. Willems, R.J.L. & van Schaik, W., (2012). Genome-wide identification of ampicillin resistance determinants in enterococcus faecium, PLoS Genet. https://doi.org/10.1371/journal.pgen.1002804.

[33] Geertsma, E.R. & Dutzler, R., (2011). A versatile and efficient high-throughput cloning tool for structural biology., Biochemistry. 50 3272–3278.

[34] Kabsch, W., (2010). XDS, 66 125–132.

[35] Schneider, T. & Sheldrick, G.M., (2002). Substructure solution with SHELXD, 58 1772–1779.

[36] Cohen, S.X. Ben Jelloul, M. Long, F. Vagin, A.A. Knipscheer, P. Lebbink, J. Sixma, T.K. Lamzin, V.S. Murshudov, G.N. & Perrakis, A., (2008). ARP/wARP and molecular replacement: the next generation, 64 49–60.

[37] McCoy, A.J., (2007). Solving structures of protein complexes by molecular replacement with Phaser, 63 32–41.

[38] Emsley, P. Lohkamp, B. Scott, W.G. & Cowtan, K., (2010). Features and development of Coot, 66 486–501.

[39] Adams, P.D. Grosse-Kunstleve, R.W. Hung, L.W. Ioerger, T.R. McCoy, A.J. Moriarty, N.W. Read, R.J. Sacchettini, J.C. Sauter, N.K. & Terwilliger, T.C., (2002). PHENIX: building new software for automated crystallographic structure determination, 58 1948–1954.

[40] Chen, V.B. Arendall, W.B. Headd, J.J. Keedy, D.A. Immormino, R.M. Kapral, G.J. Murray, L.W. Richardson, J.S. & Richardson, D.C., (2010). MolProbity: all-atom structure validation for macromolecular crystallography, 66 12–21.

[41] Waters, C.M. Hirt, H. McCormick, J.K. Schlievert, P.M. Wells, C.L. & Dunny, G.M., (2004). An amino-terminal domain of Enterococcus faecalis aggregation substance is required for aggregation, bacterial internalization by epithelial cells and binding to lipoteichoic acid., Mol. Microbiol. 52 1159–1171.

