## Supplementary information for "Enterococcal PrgA extends far outside the cell and provides surface exclusion to protect against unwanted conjugation"

[*] Shared first author

### Supplementary Tables

**Table S1.** Mass spectrometry analysis using LC-MS/MS. The numbering of the bands corresponds to the numbers on the gel in figure S1.

| Band | Ionization | Mascot Score | Mass | Num. of significant matches | Num. of significant unique sequences | Sequence coverage | Identified protein |
|---|---|---|---|---|---|---|---|
| #1 | ETD | 4097 | 143422 | 103 | 43 | 0,38 | Enterococcus faecalis PrgA |
| #2 | ETD | 7451 | 143422 | 201 | 43 | 0,38 | Enterococcus faecalis PrgA |
| #3 | ETD | 2396 | 143422 | 68 | 28 | 0,23 | Enterococcus faecalis PrgA |
| #4 | ETD | 2050 | 143422 | 71 | 31 | 0,28 | Enterococcus faecalis PrgA |
| #5 | ETD | 2731 | 143422 | 76 | 26 | 0,21 | Enterococcus faecalis PrgA |
| #6 | ETD | 2243 | 143422 | 58 | 20 | 0,15 | Enterococcus faecalis PrgA |

**Table S2**

*Enterococcus faecalis* plasmids identified to harbour both a *prgA* and a *prgB* gene. The sequence around the proposed cleavage site in PrgB is listed for each plasmid. The residues (identified in the PrgA docking studies) that are likely to interact with PrgB (PrgA1, 2 and 3, see Fig. 5) are also listed. The *prgA* and *prgB* sequences listed here can be divided into 4 distinct groups (indicated by color).

| Plasmid Accession | Plasmid | PrgB | PrgA 1 | PrgA 2 | PrgA 3 |
|---|---|---|---|---|---|
| AP018540.1 | pKUB3006-2 KUB3006 DNA | IFNYGNPKE | YMH | MNY | RLxxSHxHSxxQ |
| AP018545.1 | pKUB3007-2 KUB3007 DNA | IFNYGNPKE | YMH | MNY | RLxxSHxHSxxQ |
| CP002493.1 | EF62pB | IFNYGNPKE | YMH | MNY | RLxxSHxHSxxQ |
| NZ_CP028721.1 | pN48037F-1 | IFNYGNPKE | YMH | MNY | RLxxSHxHSxxQ |
| CP028283.1 | unnamed3 | IFNYGNPKE | YMH | MNY | RLxxSHxHSxxQ |
| AY855841.2 | pCF10 | IFNYGNPKE | YMH | MNY | RLxxSHxHSxxQ |
| AE016831.1 | pTEF2 | IFNYGNPKE | YMH | MNY | RLxxSHxHSxxQ |
| CP022485.1 | pARO1.2 | IFNYGNPKE | YMH | MNY | RLxxSHxHSxxQ |
| NZ_GG692907.1 | unnamed supercont1.10 | IFNYGNPKE | YMH | MNY | RLxxSHxHSxxQ |
| NZ_GG688649.1 | unnamed supercont1.13 | IFNYGNPKE | YMH | MNY | RLxxSHxHSxxQ |
| KT290268.1 | pPD1 | YQKKGNPKE | YGH | ENR | SSxxGHxDSxxG |
| NZ_ASWX01000006.1 | pEF10244 contig00006 | KRQFNSPKE | IGH | QNF | RMxxGHxESxxQ |
| CP028284.1 | unnamed2 | KRQFNSPKE | IGH | QNF | RMxxGHxESxxQ |
| NC_011642.1 | pMG2200 | KRQFNSPKE | IGH | QNF | RMxxGHxESxxQ |
| CP040897.1 | unnamed | KRQFNSPKE | IGH | QNF | RMxxGHxESxxQ |
| CP015885.1 | Efsorialis-p2 | KRQFNSPKE | IGH | QNF | RMxxGHxESxxQ |
| MG765452.1 | pE512 | KRQFNSPKE | IGH | QNF | RMxxGHxESxxQ |
| MK993385.1 | pEF10748 | KRQFNSPKE | IGH | QNF | RMxxGHxESxxQ |
| AP018539.1 | pKUB3006-1 KUB3006 DNA | KRQFNSPKE | IGH | QNF | RMxxGHxESxxQ |
| AP018544.1 | pKUB3007-1 KUB3007 DNA | KRQFNSPKE | IGH | QNF | RMxxGHxESxxQ |
| NZ_CP028722.1 | pN48037F-2 | KRQFNSPKE | IGH | QNF | RMxxGHxESxxQ |
| CP030044.1 | pC25-2 | YQEKGRPKQ | FDH | QNF | HSxxSHxDSxxD |
| CP042215.1 | pL15-A | YQEKGRPKQ | FDH | QNF | HSxxSHxDSxxD |
| NZ_GG670360.1 | unnamed supercont1.16 | YQEKGRPKQ | FDH | QNF | HSxxSHxDSxxD |
| CP036248.1 | pR712_02 | YQEKGRFKQ | FDH | QNF | HSxxSHxDSxxD |
| NAQY01000020.1 | unnamed1 contig_20 | YQEKGRFKQ | FDH | QNF | HSxxSHxDSxxD |
| NZ_GG692930.1 | unnamed supercont1.20 | YQEKGRPKQ | FDH | QNF | HSxxSHxDSxxD |
| CP028837.1 | unnamed2 | YQEKGRPKQ | FDH | QNF | HSxxSHxDSxxD |
| NZ_GG692709.1 | unnamed supercont1.24 | YQEKGRPKQ | FDH | QNF | HSxxSHxDSxxD |
| NC_014726.1 | pTW9 | YQEKGRPKQ | FDH | QNF | HSxxSHxDSxxD |
| CP042217.1 | pL8 | YQEKGRPKQ | FDH | QNF | HSxxSHxDSxxD |
| CP028286.1 | unnamed1 | YQEKGRPKQ | FDH | QNF | HSxxSHxDSxxD |
| CP002494.1 | EF62pC | YQEKGRFKQ | FDH | QNF | HSxxSHxDSxxD |
|  | pAD1 | YQEKGRPKQ | FDH | QNF | HSxxSHxDSxxD |
| CP019513.1 | pA | YQEKGRPKQ | FDH | QNF | HSxxSHxDSxxD |
| NZ_CP041739.1 | p1 | YQEKGRPKQ | FDH | QNF | HSxxSHxDSxxD |
| CP031029.1 | pDEF-2 | YQEKGRPKQ | FDH | QNF | HSxxSHxDSxxD |
| AE016833.1 | pTEF1 | YQEKGRPKQ | FDH | QNF | HSxxSHxDSxxD |

### Supplementary Figures

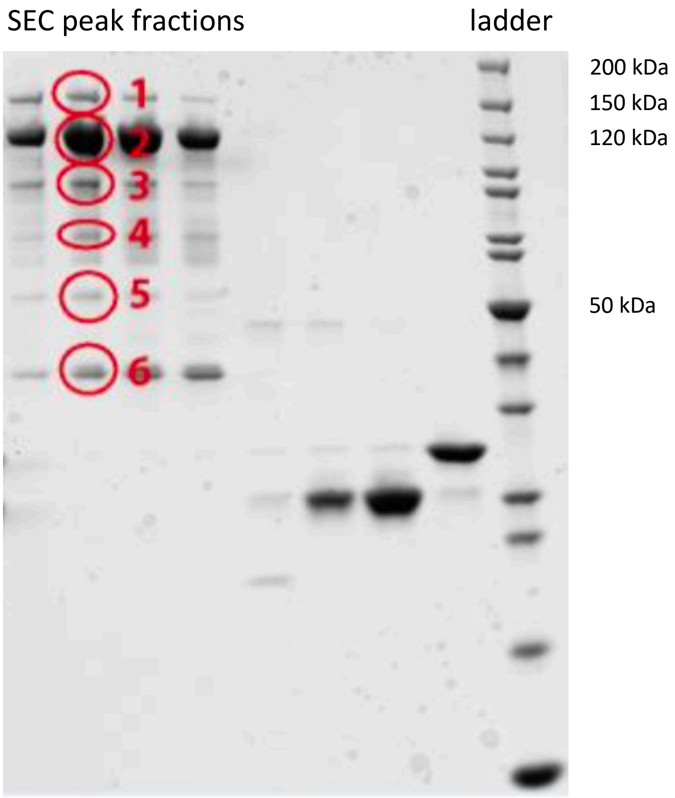


**Supplementary Figure 1.** SDS-PAGE of the SEC peak fractions from a His-PrgA$_{28\text{-}814}$ purification. The bands highlighted in red circles came from the main gelfiltration peak and were analysed via mass spectrometry. All bands were identified to originate from PrgA$_{28\text{-}814}$, but only a small fraction of the purified protein was full-length (band #1).

```
CLUSTAL O(1.2.4) multiple sequence alignment


PrgA-CAP-domain     ------------------------------------------------------------ 0
5JYS                KLSKLSILTSALATSALAAPAVVTVTEHAHEAAVVTVQGVVYVENGQTRTTYETLAPAST 60
1U53                ------------------------------------------------------------ 0
3Q2R                ------------------------------------------------------------ 0
1XTA                ------------------------------------------------------------ 0
2DDA                ------------------------------------------------------------ 0


PrgA-CAP-domain     --------------------------------------------QNTLDNSK---EELKG 303
5JYS                ATPTSTATALVAPPVAPSSASSNSDVVLSALKNLASVWGKTTDSTTTLTSSESTSQSLAQ 120
1U53                ------------------------------------------------------------ 0
3Q2R                ------------------------------------------------------------ 0
1XTA                ------------------------------------------------------------ 0
2DDA                ------------------------------------------------------------ 0


PrgA-CAP-domain     HKGINLPPKFSADYDTKLSAEEIATLEKTALEMNKNFPTSKEDEKNKDVMWDIQHLSADQ 363
5JYS                ATTTSTPAAAS--------TTSTPAATTTTSQAAATSSASSSDSDLSDFASSV------- 165
1U53                --------------------------------AEAEFGCPDNG-MSEEARQKF------- 20
3Q2R                -------------------------------EAEAEFANILPDIENEDFIKDC------- 22
1XTA                ---------------------------------NVDFN--SESTRRKKKQKEI------- 18
2DDA                -------------------------------------------SNKKNYQKEI------- 10
                                                                  ..   .

PrgA-CAP-domain     KKELSVYTTELLNDVRKKLGLSQL--------SVSDQSIKFAWDIAKYSDTGEYMHDVIA 416
5JYS                --------LAEHNKKRAL----------------HKDTPALSWSDTLASYAQDYADNYDC 201
1U53                --------LEMHNSLRSSVALGQAKDGAGGNAPKAAKMKTMAYDCEVEKTAMNNAKQCVF 72
3Q2R                --------VRIHNKFRSEV------------KPTASDMLYMTWDPALAQIAKAWASNCQF 62
1XTA                --------VDLHNSLRRRV------------SPTASNMLKMEWYPEAASNAERWANTCSL 58
2DDA                --------VDKHNALRRSV------------KPTARNMLQMKWNSRAAQNAKRWANRCTF 50
                                *  *                    .   : :     . :

PrgA-CAP-domain     IN-----------------------------------KAAKENGFKEYPGMNYYE--NLG 439
5JYS                SG--------TLTHSGGPYGENLALGYD--------GPAAVDAWYNEISNYDFSN----- 240
1U53                KHSQPN--------QRKGLGENIFMSSDSGMDKAKAAEQASKAWFGELAEKGVGQNLKLT 124
3Q2R                SHNTRLKPPHKLHPNFTSLGENIWTGSVPIFS----VSSAITNWYDEIQDYDFKT--RIC 116
1XTA                NHSPD----NLRVLEGIQCGESIYMSSNA-RT----WTEIIHLWHDEYKNFVYGVGASPP 109
2DDA                AHSPP----NKRTVGKLRCGENIFMSSQP-FP----WSGVVQAWYDEIKNFVYGIGAKPP 101
                                                                . *

PrgA-CAP-domain     GGYYETENGKVSKYTLQESIRKMLVNMLFDDGRLGYSHLHSLLQDGKTALGVSLSGEKNS 498
5JYS                -PGFSSNTGHFTQVVWK----------------------------STTQVGCGIKTCGGA 271
1U53                GGLFSRGVGHYTQMVWQ----------------------------ETVKLGCYVEACSNM 156
3Q2R                ----KKVCGHYTQVVWA----------------------------DSYKVGCAVQFCPKV 144
1XTA                ----GSVTGHYTQIVWY----------------------------QTYRAGCAVSYCPSS 137
2DDA                ----GSVIGHYTQVVWY----------------------------KSYLIGCASAKCSSS 129
                            *: :: .                               .   *

PrgA-CAP-domain     -----I-SPKIHIISYGKEK---LEDSSQYQNGEVASMKSK-EELQQEIASNQ------- 541
5JYS                -------WGDYVICSYDPAGNYEGEYADNVEPLA-------------------------- 298
1U53                ---------CYVVCQYGPAGNMMGKD--IYEKGEPCSKCENCDK-EKGLCSA-------- 196
3Q2R                SGFDALSNGAHFICNYGPGGNYP---TWPYKRGATCSACPNNDKCLDNLCVNRQRDQVK- 200
1XTA                A------WSYFYVCQYCPSGNFQGKTATPYKLGPPCGDCPS--ACDNGLCTNPCTIYNKL 189
2DDA                --------KYLYVCQYCPAGNIRGSIATPYKSGPPCADCPS--ACVNKLCTNPCKRNNDF 179
                                : .*              :

PrgA-CAP-domain     --------------------------------       541
5JYS                --------------------------------       298
1U53                --------------------------------       196
3Q2R                -RYYSV--------------------------       205
1XTA                TNCDSLLKQSSCQDDWIKSNCPASCFCRNKII       221
2DDA                SNCKSLAKKSKCQTEWIKKKCPASCFCHNKII       211
```

**Supplementary Figure 2.** Sequence alignment of the CAP domain of PrgA with the CAP domains of other structurally characterized proteins. Although the structures are very similar (Fig. 1), the proteins have very low sequence identity.

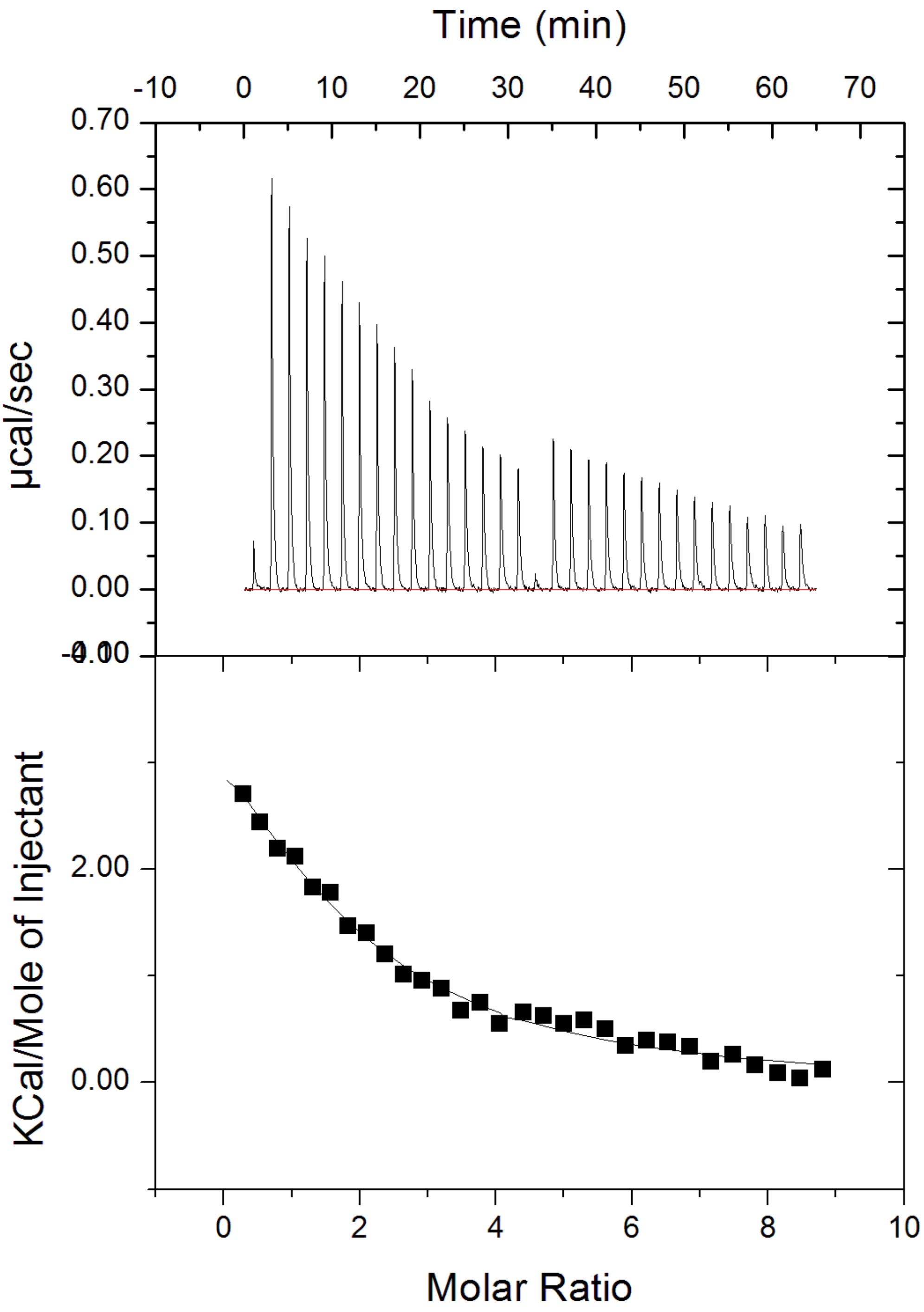


**Supplementary Figure 3.** ITC of PrgA titrated with $ZnSO_4$. To obtain a full titration curve two consecutive titrations were done into the same protein sample, and the resulting curves merged into one. The 2nd titration starts with the low peak in the middle of the upper graph.

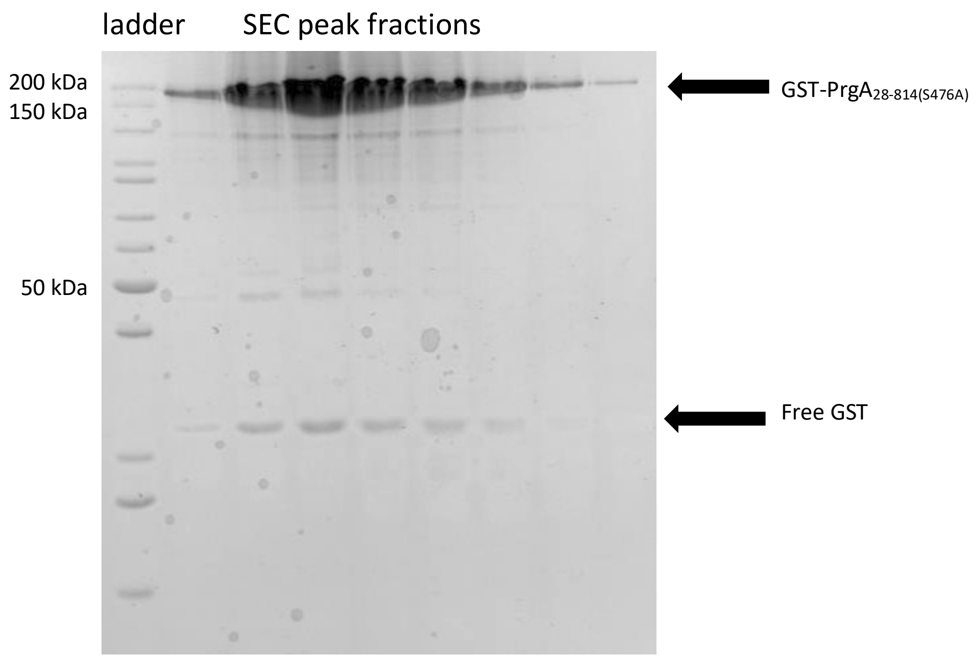


**Supplementary Figure 4**. SDS-PAGE of purification of GST-PrgA$_{28\text{-}814(S476A)}$. As compared to Fig S1, it´s evident that the large majority of GST-PrgA$_{28\text{-}814(S476A)}$ is in its full-length form, while in Fig S1 (where the wild-type protein was purified without the addition of divalent cations) only a small fraction is full-length.

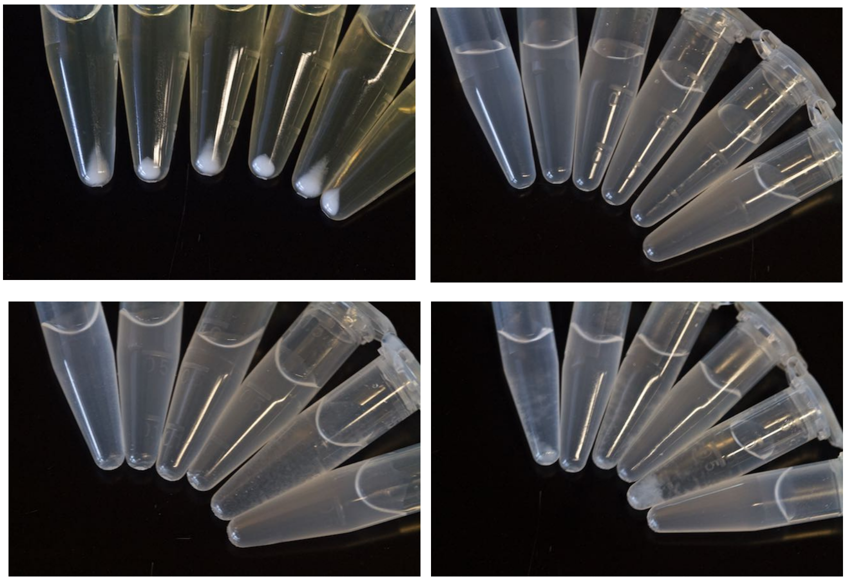

**Supplementary figure 5.** Depiction of aggregate formation assay. (Upper left) Pelleted cells. (Upper right) the same cells after resuspension in PUM-buffer, (Lower left) immediately after vortexing and then (Lower right) after 15 min settling. The tubes from left to right in each picture are pCF10: first induced, then uninduced, pCF10:prgA:S476A induced/uninduced and pCF10:ΔprgA induced/uninduced. oD of supernatant is determined after 15 min of settling and compared to the uninduced culture. Small aggregates are visible in the induced pCF10:ΔprgA immediately after vortexing (C, 2nd tube from the right).

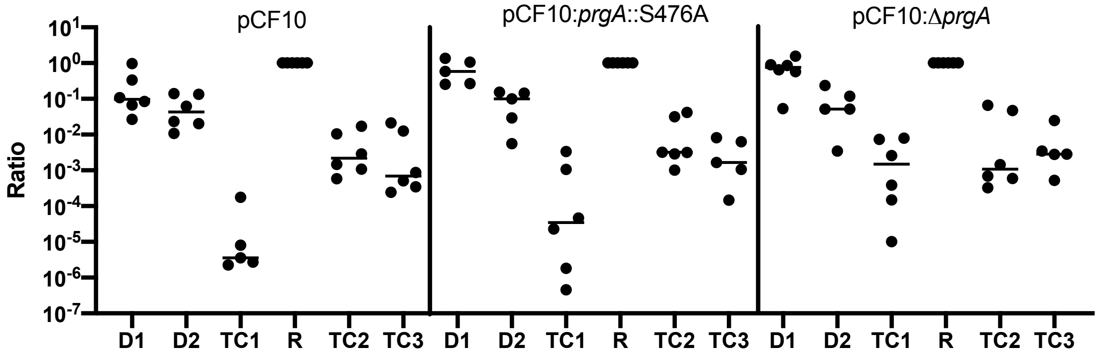


**Supplementary figure 6.** Population Analysis in a three partner mating. Recipient strain OG1ES (R) was mixed with OG1Sp:pCF10-G2 (D1) and OG1RF:pCF10 (wild-type or variants, D2). Cells were incubated on an agar surface for three days and then enumerated. TC1: Transconjugants resulting from the transfer of pCF10-G2 (from D1) into D2 strains. TC2 and TC3 represent transconjugants into the recipient strain OG1ES. Cell numbers were normalized to the cell number of the recipients in each experiment (R=1).

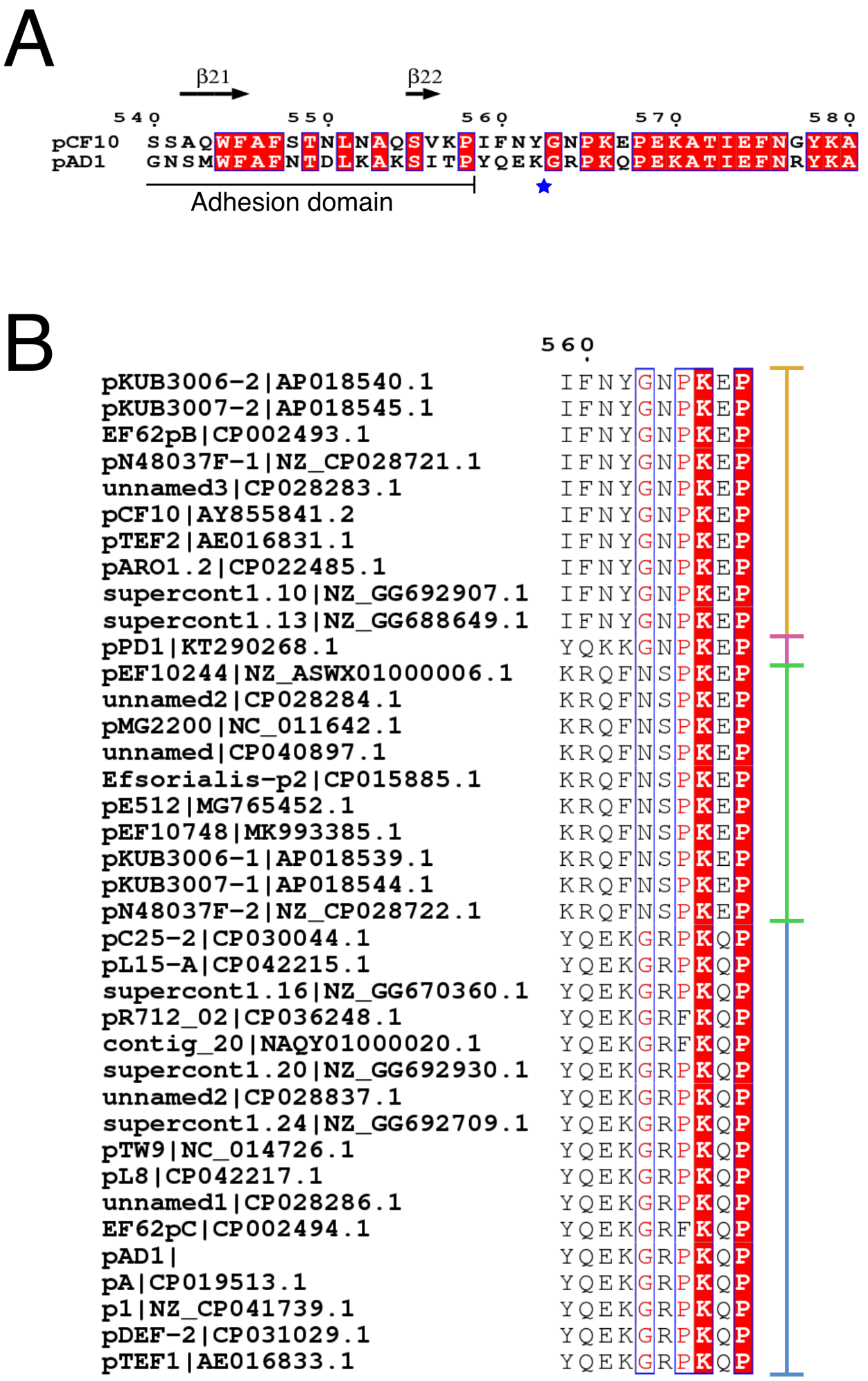


**Supplementary Figure 7.** Sequence analysis of *prgB* showing potential cleavage sites and groupings. A: Sequence alignment of *prgB* from pCF10 with *prgB* from pAD1 depicting the region around the proteolytic cleavage site in $prgB_{\mathrm{pAD1}}$ (marked by a blue star). The secondary structure elements of the N-terminal adhesion domain are depicted above the sequence and the end of the domain is marked below. The region downstream of the adhesion domain is predicted to be disordered. B: Sequence alignment of the proposed proteolytic cleavage site in *prgB* genes in plasmids that also contain a *prgA* gene. These sequences can be placed in 4 groups as highlighted by coloured lines to the right of the alignment, also shown in Table S1.

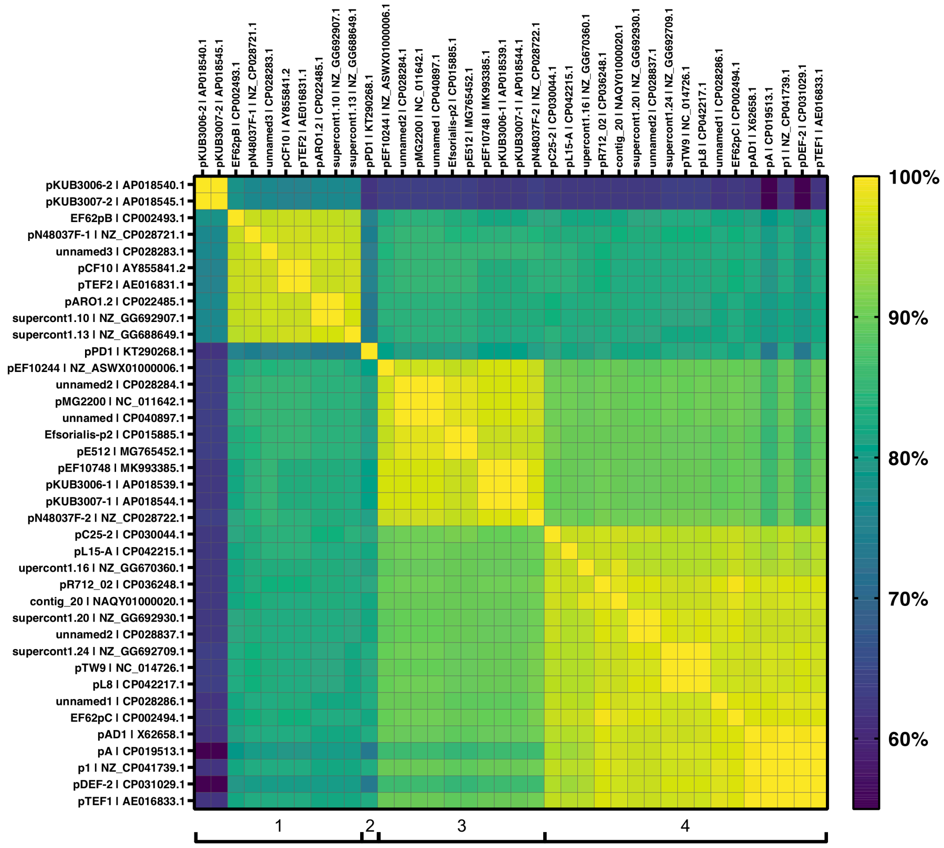


**Supplementary Figure 8.** Percent sequence identity matrix comparing the *prgA* genes from Enterococcus plasmids annotated in the NCBI database that also contain a *prgB* gene. *prgA* genes cluster into four distinct groups. These groups coincide with the *prgB* clusters (see Fig. S7B and Table S1).
